# A unified framework for hydromechanical signaling: Do plant signals go with the flow?

**DOI:** 10.1101/2024.10.23.619842

**Authors:** Vesna Bacheva, Fulton Rockwell, Jean-Baptiste Salmon, Jesse Woodson, Margaret Frank, Abraham Stroock

**Affiliations:** Smith School of Chemical and Biomolecular Engineering, Cornell University, Ithaca 14853, NY; School of Integrative Plant Science, Cornell University, Ithaca, NY 14853, USA; Kavli Institute at Cornell for Nanoscale Science, Cornell University, Ithaca, NY 14853, USA; Department of Organismal and Evolutionary Biology, Harvard University, Cambridge, MA 02138, USA; CNRS, Syensqo, LOF, UMR 5258, Univ. Bordeaux, F-33600 Pessac, France; The School of Plant Sciences, University of Arizona, Tucson, AZ 85721, USA

## Abstract

Local wounding in plants triggers signals that travel locally within the wounded leaf or systemically through the vasculature to distal leaves. The transmission mechanisms of this ubiquitous class of signals remain poorly understood. Here, we develop a unifying framework based on poroelastic dynamics to study two coupled biophysical processes – propagation of pressure changes and transmission of chemical elicitors via mass flows driven by these pressure changes – as potential mechanisms for the initiation and propagation of wound-induced signals. We show that rapid pressure changes in the xylem can transmit mechanical information across the plant, while their coupling with neighboring non-vascular tissue drives swelling and mass flow that can transport chemical elicitors to distal leaves. We confront our model predictions with signaling dynamics measurements in several species, and show that the poroelastic model captures observed mechanical, biochemical, and electrophysiological signals. This framework provides a valuable foundation for assessing mechanisms of signal transmission and for designing future experiments to elucidate factors involved in signal initiation, propagation, and target elicitation.

## Introduction

As sessile organisms, plants are remarkably adaptative to abiotic and biotic perturbations imposed by their local environments.^1^ While these perturbations are often localized within a tissue, plants can transduce local and abrupt stimuli into whole-organism responses via relatively fast (seconds to minutes) transmission of chemical, electrical, and mechanical signals.^2^ In the context of wound-induced perturbations, such as by burning, crushing, or cutting, studies dating back more than a century have proposed and elucidated the propagation of specific chemical elicitors,^3–5^ changes in turgor pressure,^6,7^ tissue deformation,^8^ and propagation of slow wave potentials (Fig. 1a).^9^ Important questions remain regarding the details of these signals, including the mechanisms of their upstream initiation and downstream transduction, the coupling among biochemical and biophysical processes involved in propagation, and their dependence on specific plant physiological traits and states. This incomplete understanding hinders the development of predictive models for plant stress responses and efforts to enhance crop management and performance through breeding and bioengineering.

**Fig. 1.**
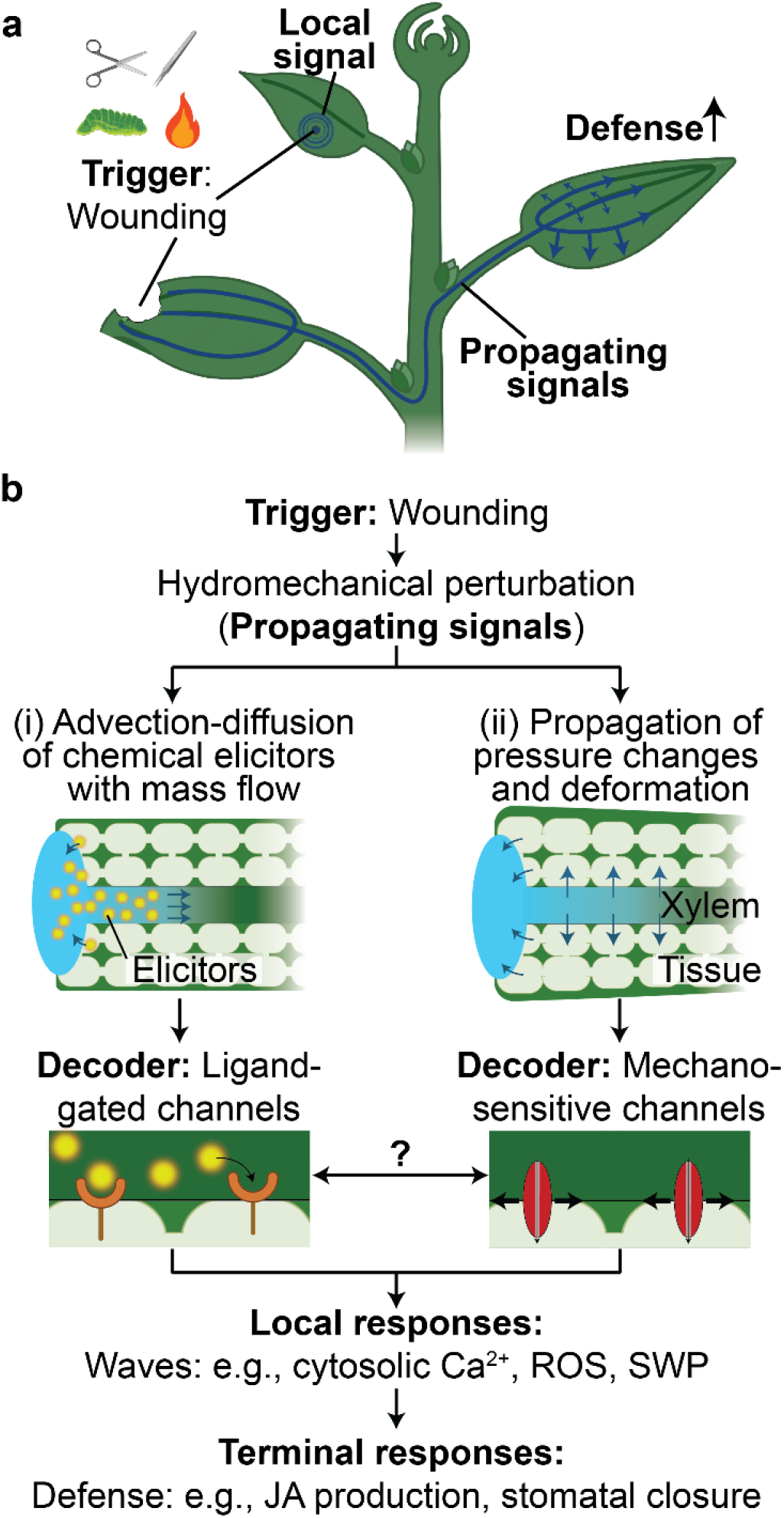
Wound-induced signaling in plants. **a**, Local wounding (e.g., burning, crushing, cutting, herbivore attack) triggers propagating signals that can travel locally in the damaged leaf or via the vasculature to distal leaves, to induce systemic defense responses. **b**, A local wound triggers hydromechanical perturbations with the release of symplastic water into the apoplast. This release of water initiates two coupled physical processes that can serve to propagate these biological signals: (i) mass flow (motion of water - left) that can carry chemical elicitors (i.e., Ricca factors) by advection and diffusion to downstream sites of interaction, for example by ligand-gated channels or receptors; (ii) Propagation of the pressure changes (relaxation of tension - right) accompanied by tissue deformation (dilation). These changes in stress and strain in the tissue may act directly as propagating signals that can be decoded mechanically, for example by mechanosensitive ion channels. These two decoding pathways may be coupled and trigger different local responses (e.g., cytosolic calcium (Ca^2+^), reactive oxygen species (ROS), slow wave potential (SWP) waves) that participate in the propagation processes and mediate terminal defense responses such as production of the wound hormone jasmonic acid (JA) and closure of stomates.

The mechanisms underlying signal propagation have been the subject of much debate. One hypothesis,^4,5,10,11^ dating back to Ricca in 1916,^3^ proposes that a chemical elicitor, termed ‘Ricca factor’, released at the wound site and propagated via advection and diffusion in the vasculature with mass flow (i.e., motion of water), activates ligand-gated channels in neighboring cells (Fig. 1b(i)). Several elicitors have been suggested as Ricca factors, including amino acids, e.g., glutamate^5,10^ and enzymes, e.g., thioglucosidase.^12^ Another hypothesis suggests that the propagation of hydraulic waves (i.e., pressure changes) along the vasculature serve as both the mode of transmission and the signal,^6,7,13–15^ with propagating changes in pressure triggering local pathways mediated by mechanosensitive ion channels (Fig. 1b(ii)).^16^ Alternatively, both advective processes (i.e., transmission of chemical elicitors) and mechanical processes (i.e., propagation of pressure changes) may work together for signal propagation.^16^

Among the potential mechanisms for the initiation and propagation of wound-induced signals, hydromechanical processes - involving water flow and transmission of mechanical stresses and strains through plant tissues - stand out because they are inevitably triggered at a site of wounding due to the disruption of the native pre-stresses associated with the imbalance of pressures between apoplast, symplast, and the atmosphere in turgid tissues. These hydromechanical processes can be described with poroelasticity, a modeling framework that accounts for plant structure formed of porous and elastic materials filled with water.^17–21^ The mode of propagation of hydromechanical processes favors vascular pathways, through which the propagation of various systemic signals have been observed, for example, with gene-encoded reporters of cytosolic calcium concetration.^5,10,16^ Poroelastic transmission of fluid flow and stresses also operates in non-vascular tissues in response to local (e.g., cell-scale manipulations)^22,23^ and global (e.g., changes in evaporative demand)^17,24^ perturbations. While roles for hydromechanical processes have frequently been hypothesized in the initiation and propagation of responses to wounding events (Fig. 1b),^4,11,25–27^ previous models of these processes have not addressed the coupling of xylem and tissue, whole-plant architecture, or mechanistic bases for the dynamics of the underlying stresses and mass flows (see S.I. Section S1). A predictive model that captures these important features could provide a biophysical foundation for the design and interpretation of experiments aimed at decoding these critical plant pathways.

Here, we derive a unifying framework based on poroelastic dynamics that couples mechanical and advective processes resulting from hydromechanical perturbations in a wounded plant and use this model to study how these processes could carry biological information. Compared to existing models, our framework: (i) integrates both advective and mechanical processes with explicit coupling between the xylem and non-vascular tissues, (ii) accounts for the architecture of the entire plant, including roots, stems, and leaves, to predict the propagation of both systemic and local signals; (iii) offers a biophysically grounded explanation for the underlying mass flows; and (iv) defines all model inputs using physiologically relevant and experimentally measured parameters. In the following, we show that, with the unification of these features, poroelastic dynamics can explain observations of mechanical,^8^ biochemical,^5^ and electrophysiological signals,^27^ both qualitatively and quantitatively, and clarify important outstanding questions.

## Results

### Plant tissues as poroelastic media

Many biological tissues, including those of plants can be viewed as a porous, permeable, and deformable media filled with water.^17–21^ The water in plants can move through apoplastic (i.e., via xylem and cell wall), symplastic (i.e., via plasmodesmata (PD)) and cross-membrane (i.e, plasma-membrane mediated) pathways. The water through the symplastic and apoplastic pathways occurs along gradients of pressure, *P* [MPa], whereas the water through the cross-membrane occurs through gradient of water potentials, *ψ* [MPa], due to the presence of semi-permeable membranes (See S.I. Section S2.A). The poroelastic theory initially formulated by Biot^28^ in the context of soil mechanics, and later extended to plant cells by Philip,^18^ has been successfully adapted to predict the dynamics of mechanical responses to perturbations in water potential in plant leaves^17^ and branches^20,29^. However, these studies have considered neither the whole-plant architecture nor the coupling between xylem and non-vascular tissues, and they have not been adapted to explore wound-induced signals and responses (See S.I. Section S1.A).

Fig. 2a illustrates our poroelastic model of a wounded plant. Here, we focus on the specific assumptions outlined in the following sentences (see also S.I. Section S2.A); in the S.I. we provide a discussion of generalization of the framework beyond these assumptions (see S.I. Section S3). We model plants as poroelastic media composed of two coupled compartments: a xylem bundle composed of individual xylem vessels and living cells that we refer to as non-vascular tissue. The water flow through the xylem occurs through apoplastic paths; water flow in the tissue occurs through parallel symplastic and cross-membrane paths. We treat the tissue as an effective material in which the apoplast and symplast are in local equilibrium (See S.I. Section S2.A), and we assume that during the relatively fast (seconds to minutes) propagation of wound-induced responses, the tissue osmotic pressure remains constant and uniform such that water flow in the tissue occurs along gradients of turgor pressure. We focus our analysis on two leaves that share direct vascular connections. The leaves span a distance, *L* [m], the spacing between the leaf veins is 2*W* [m], and the leaf thickness is *d*_*th*_ [m]. The leaves transpire from their top and bottom surfaces with a constant transpiration flux, *E* [mol m^-2^ s^-1^]. We represent the node where the two leaves are connected to the stem as a water source that replenishes the water lost through transpiration. Our explicit inclusion of xylem architecture and xylem-tissue coupling provides an opportunity to examine the dynamics of scenarios involved in signaling.

**Fig. 2.**
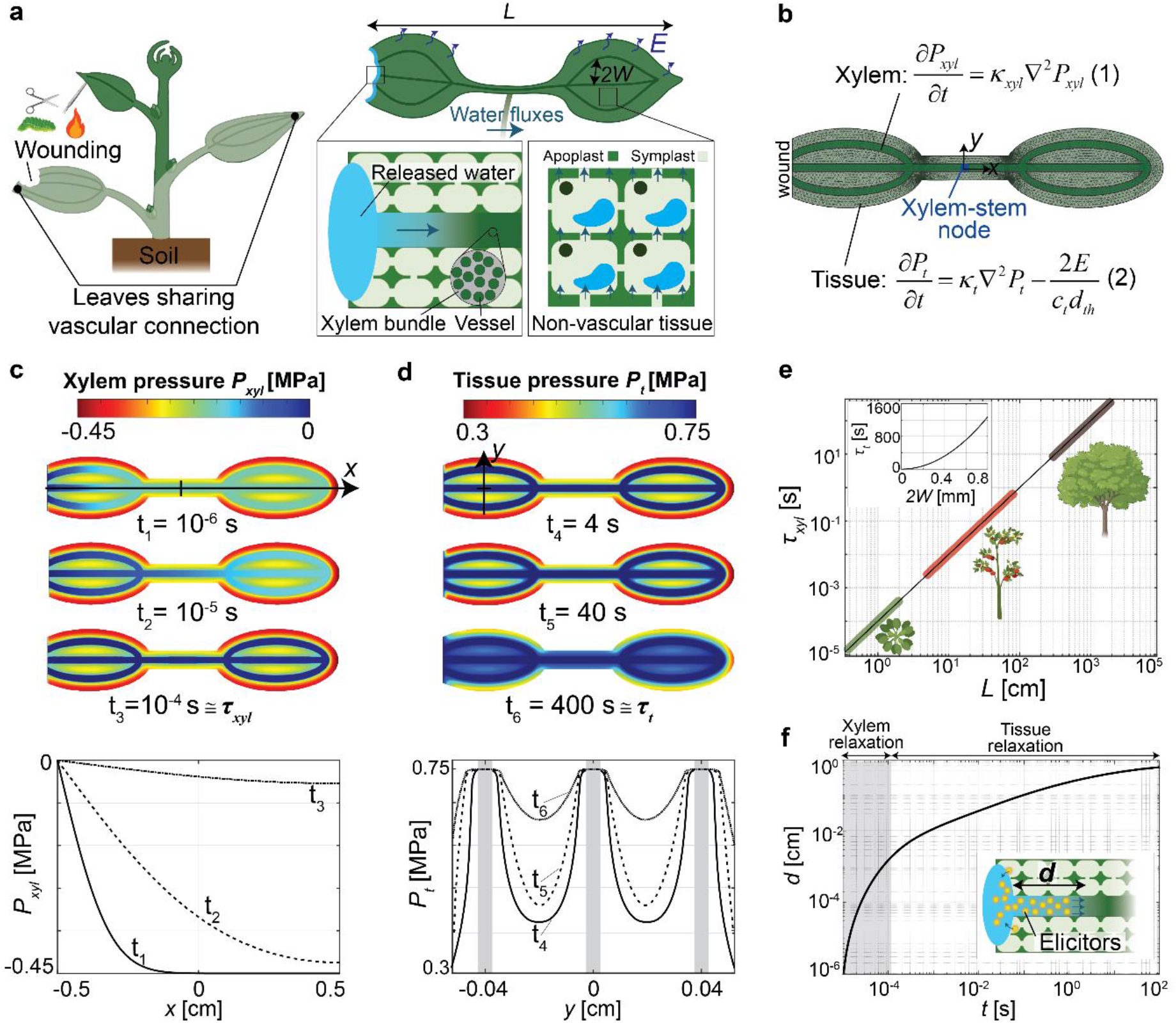
Poroelastic framework and predicted responses. **a**, Schematic representation of a wounded plant, focusing on a distal leaf that shares a direct vascular connection with the wounded leaf. The leaves span a distance *L* [m], the spacing between the leaf veins is 2*W* [m], and the leaf thickness is *d*_*th*_ [m]. The leaves transpire from their top and bottom surfaces with a constant transpiration flux, *E* [mol m^-2^ s^-1^]. Following wounding, ruptured cells release water at the wound site which propagates via the xylem bundle, and from the xylem to non-vascular tissue. We assume that this released water is not depleted during the simulated relaxations, and we define this as a “wet wound”. **b**, Meshed representation of the wounded plant used to numerically solve the resulting gradients of xylem and tissue turgor pressure, *P* [MPa]. We represent the node where the two leaves are connected to the stem as a water source that replenishes the water lost through transpiration. Upon wounding, the pressure in the xylem (*P*_*xyl*_) and tissue (*P*_*t*_) will shift towards a new equilibrium. These processes are governed by poroelastic diffusion equations for xylem (Eq. 1) and tissue (Eq. 2). This diffusive dynamic of the pressure is characterized by the poroelastic diffusivities of the xylem, *κ*_*x*_ [m^2^s^-1^], and the tissue, *κ*_*t*_. **c-d**, Numerical predictions for the evolution of pressure in the xylem (c) and tissue (d) based on Eqs. 1 and 2 (b) in a small herbaceous plant (e.g., *Arabidopsis thaliana*) with a leaf-to-leaf distance of *L =* 1 cm and vein spacing of 2*W =* 0.4 mm. Upon wounding, the released water and tension propagates rapidly through the xylem such that the pressure along this path approaches equilibrium (dark blue) within a relaxation timescale, *τ*_*xyl*_ = *L*^*2*^*/ κ*_*xyl*_ ≅ 100 μs. The water efflux from the xylem along its entire length into the tissue induces an increase of tissue pressure over time. The tissue pressure approaches equilibrium within its relaxation timescale, *τ*_*t*_*=W*^*2*^*/κ*_*t*_ ≅ 400 s. The plots below show the variation of the xylem pressure in the middle vein (along *x* in (c)), and the tissue pressure across the leaf (along *y* in (c)) for the three different times. Time-lapse videos of the propagation of pressure changes in the xylem and tissue are provided in Movie S2 and S3. **e**, Plot of the predicted xylem relaxation timescale for plants of different sizes. Since the timescale for relaxation of the xylem varies quadratically with the length of the plant (*τ*_*xyl*_*∼L*^*2*^), our model predicts that larger plants will exhibit a significantly longer relaxation time. The inset shows the predicted relaxation time in the tissue which varies quadratically with the vein spacing (*τ*_*t*_*∼W*^*2*^). **f**, Numerical prediction of the distance chemical elicitors (i.e., Ricca factors) released at the wound site propagates in the xylem by advection and diffusion (molecular diffusivity, *D* = 10^−10^ m^2^s^-1^). Displacement with the rapid xylem pressure equilibration occurs rapidly (within 100 μs) and is small compared to the dimension of the plant (*d* = 20 μm << *L*=1 cm); the continued displacement with the flow driven by transpiration and tissue relaxation is slower (hundreds of seconds) and can reach the dimension of the plant (*d* = 0.8 cm ≅ *L*). All numerical predictions shown in this figure (**c, d, f**) were simulated for a transpiring herbaceous plant (e.g., *Arabidopsis thaliana*) with initial distribution of xylem and tissue turgor pressure shown in Fig. S2, and using experimentally available parameters: *L* = 1 cm, 2*W* = 0.4 mm, *κ*_*xyl*_ =1 m^2^s^-1^, *κ*_*t*_ *=*10^−10^ m^2^s^-1^, *P*_*0*_ *=* -0.2 MPa, *E* = 1 mmol m^-2^ s^-1^, *c*_*t*_ = 10^−2^ mol m^-3^ Pa^-1^, *d*_th_ = 200 μm, *D* = 10^−10^ m^2^s^-1^, *π*_*t*_ = 0.75 MPa, *h*_*stem*_ = 10^−8^ mol m^-2^ Pa^-1^ s^-1^ (See S.I. Sections S5 and S7 and Table S2).

Wounding events locally disrupt the mechanical integrity of the cell membrane and release water previously constrained within the symplast (Fig. 2a – expanded view of wound). As water within the xylem is typically under tension (negative pressure), the available water at the wound site is pulled into the xylem, and the local tension in the xylem is released. The water then moves from the xylem to non-vascular tissue. It is important to note that while our analysis here focuses on wounding events involving water release due to mechanical rupture of cell membranes, our model is also appropriate for responses to non-wounding events, such as touching and squeezing in which passive or active release water from the symplast into the apoplast could occur, as observed in rapid leaf folding in *Mimosa pudica*.^30^

Fig. 2b shows the meshed representation of the wounded plant and the governing equations that we use to solve numerically for the propagation of changes in xylem and tissue turgor pressure, and the resulting water mass flows. Important for the propagation of pressures changes, and therefore other downstream processes, is that poroelastic transients are intrinsically diffusive in nature as governed by the poroelastic diffusion equations for the xylem (Eq. 1) and tissue (Eq. 2) shown in Fig. 2b (See S.I. Section S2 for model derivation). Analogous to molecular diffusion, this diffusive dynamic of the pressure is characterized by the poroelastic diffusivities of the xylem, *κ*_*xyl*_ [m^2^s^-1^], and that of the surrounding tissue, *κ*_*t*_ [m^2^s^-1^]. The poroelastic diffusivity is defined by the ratio of the hydraulic conductivity (*k*_*xyl*_, *k*_*t*_ [mol m^-1^ Pa^-1^ s^-1^] – how easily water moves through xylem and tissue), and the hydraulic capacity (*c*_*xyl*_, *c*_*t*_ [mol m^-3^ Pa^-1^] – how much the xylem and tissue dilate with changes in pressure): *κ*_*xyl*_ *= k*_*xyl*_ /*c*_*xyl*_, *κ*_*t*_ *= k*_*t*_ /*c*_*t*_. The xylem vessels have relatively low resistance to water flow (large *k*_*xyl*_), and their reinforced cell walls make them relatively inelastic (low *c*_*xyl*_),^6^ resulting in large poroelastic diffusivity (*κ*_*xyl*_ ∼ 1 m^2^ s^-1^) relative to the typical molecular diffusivity of a small molecule (*D* ∼ 10^−10^ m^2^ s^-1^).^31^ In contrast, surrounding tissue is composed of a network of relatively low-conductance and highly elastic cells, resulting in a relatively low poroelastic diffusivity (*κ*_*t*_ ∼ 10^−10^ m^2^ s^-1^) compared to that in the xylem.

Fig. 2c-d present snapshots of the pressure field from a simulation (top - see Movies S2 and S3) with profiles of the pressure along the xylem (bottom, Fig. 2c) and tissue (bottom, Fig. 2d) for a small herbaceous (non-woody) plant (e.g., *Arabidopsis thaliana*) with a leaf-to-leaf distance of *L =* 1 cm and vein spacing of 2*W =* 0.4 mm. In Fig. 2c, we resolve the rapid transient of pressure in the xylem characterized by a short relaxation time, *τ*_*xyl*_ [s] = *L*^2^/*κ*_*xyl*_ ≅ 100 μs (time at which pressure throughout vessels becomes blue, nearly 0 MPa). The plot in Fig. 2e shows the predicted growth of *τ*_*xyl*_ with the global dimension of plants, *L* (see S.I. Section S2.B for scaling analysis for *τ*_*xyl*_). In Fig. 2d, we resolve the slower transient of pressure in the adjacent tissue characterized by a relaxation time, *τ*_*t*_ *=W*^2^/*κ*_*t*_ ≅ 400 s (time at which pressure throughout tissue becomes blue, the maximum turgor pressure of 0.75 MPa). The rapid propagation of the change in xylem pressure initiates the transient in the tissue nearly simultaneously throughout the plant. The inset plot in Fig. 2e shows the growth of *τ*_*t*_ over the typical range in spacing between veins across species (see S.I. Section S2.B for scaling of *τ*_*t*_). Unlike the variation in plant size, the spacing of leaf veins is relatively conserved (2*W* ∼ 0.1-1 mm), suggesting that the tissue relaxation timescale will be similar across different species. In Fig. 4f, we report the predicted distance, *d* [cm] that a chemical elicitor would be advected by the mass flows in the xylem induced by the relaxation of the xylem itself (grey area) and by the subsequent relaxation of the surrounding tissue (which pulls additional water through the xylem).

These predictions inform the following insights about hydraulic transients triggered by wounding: (i) For small plants (*L* < 100 cm), the propagation of changes of pressure through the xylem after wounding is fast (*τ*_*xyl*_ < 1 s – Fig. 2e) relative to both the time over which tissues relax (*τ*_*t*_ =10-10^3^ s – Fig. 2e inset) and the time scale on which systemic wounding responses are typically observed (10-100 s). This separation of time scales suggests that, while xylem can rapidly communicate mechanical information about a wounding event to all tissues, it is unlikely to be directly and independently responsible for the systemic signals (e.g., calcium or electrical) that have been reported to date. Alternatively, it may directly trigger responses that have not been observable with the reporters used to date or due to insufficient temporal resolution of the measurement techniques employed (see S.I. Section S9); (ii) The maximum predicted distance for the advection of an elicitor factor based on the mass flow generated by relaxation of the xylem (*d* ≅ 10^−3^ cm – Fig. 2f) is orders of magnitude smaller than required to mediate long-distance signaling; (iii) The range of times predicted for the relaxation of tissue (10-10^3^ s – Fig. 2e inset) is commensurate with times over which long-distance signals have been characterized and the predicted distance of advection in xylem flows generated by this process (*d* ≅ 1 cm – Fig. 2f) is compatible with observed whole-plant signals. Based on these points, we pursue our analysis of long-distance signals with a focus on the processes mediated by the relaxation of tissues (Fig. 2d) that we expect to be initiated nearly simultaneously throughout after wounding by the rapid relaxation of xylem (Fig. 2c).

### Wound-induced leaf swelling follows poroelastic dynamics

We begin by confronting the predictions of our poroelastic model with observations of purely mechanical, systemic responses in tissues driven by wounding. In experiments with various plants, Malone demonstrated that localized scorching of one leaf results in an increase in turgor pressure^13^ and swelling^6,8,31^ of a neighboring leaf. Recent studies have also reported that wounding can lead to petiole bending^15,32^ and leaf movement.^33,34^ Fig. 3a illustrates experimental observations by Malone of changes in relative leaf thickness in wheat seedlings measured by transducers placed at five different locations along a leaf neighboring a scorched leaf.^8^ We note two features in Malone’s observations: (i) the increase in leaf thickness appeared to start simultaneously at all positions along the leaf after the wounding event (arrow with flame); and, (ii) following this initiation, the dynamics of the changes in leaf thickness varied as a function of position along the length (T0-T4), with, counter intuitively, equilibration occurring more quickly for more distal positions (e.g., T4) than for more proximal ones (e.g., T0) along the leaf.

**Fig. 3.**
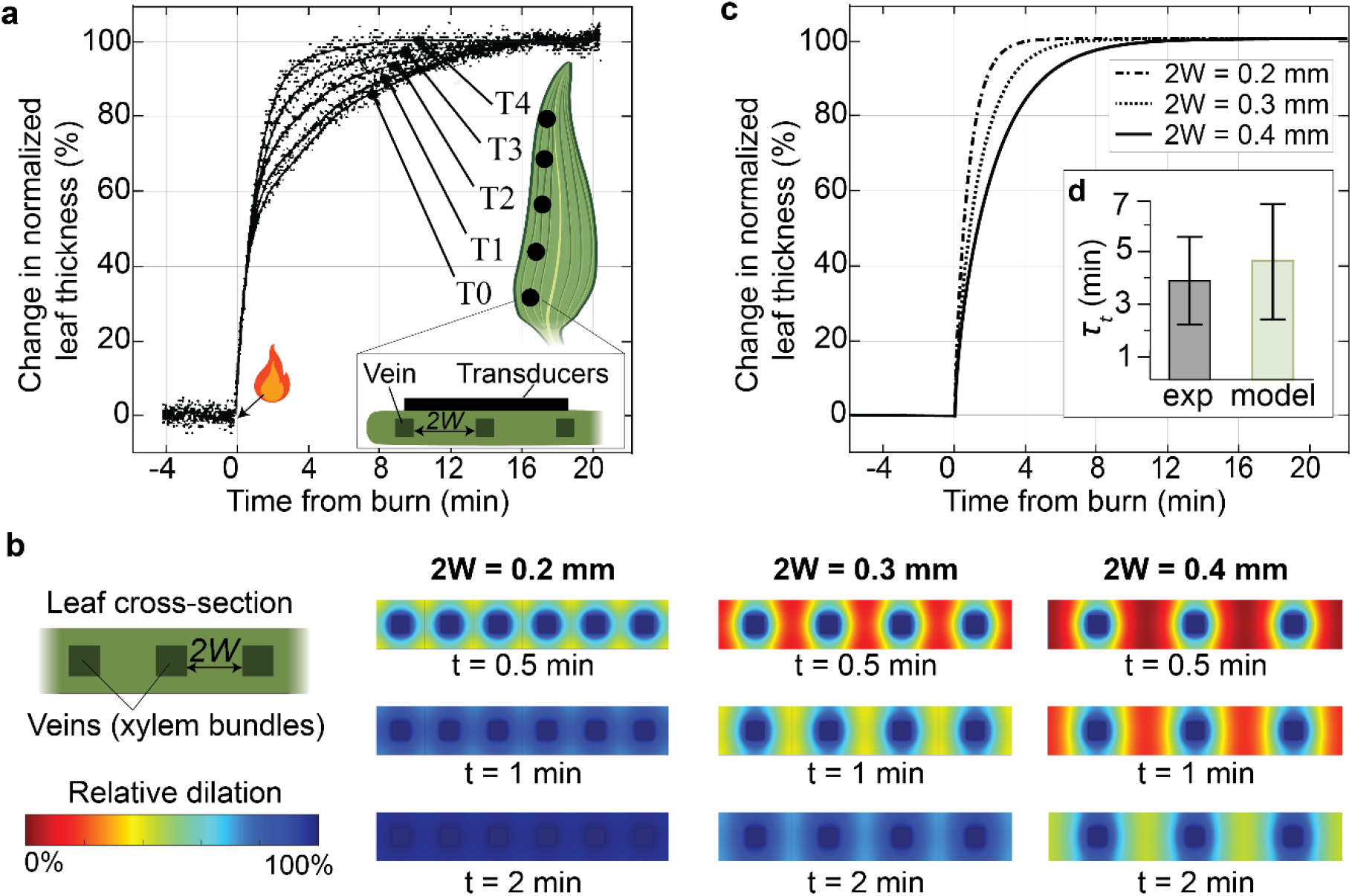
Long-distance propagation of wound-induced leaf deformation. Experimental measurements obtained by Malone^8^ in wheat of the kinetics of the increase in the leaf thickness as measured with displacement transducers at five positions (T0, T1, T2, T3, T4) along a leaf neighboring a leaf wounded by scorching. The vein-spacing (2*W* [m]) decreases along the length of the leaf, towards the tip. The plot shows the change in normalized leaf thickness expressed relative to the maximum change in leaf thickness at a given location. Malone^8^ observed that the increase in leaf thickness started simultaneously across at all positions along the leaf (*t* = 0). The overlapping initial transient at all locations suggests rapid equilibration of the xylem pressure, consistent with the prediction of the rapid xylem relaxation timescale from our poroelastic model (Fig. 2c). **b**, Numerical prediction of the relative leaf dilation (local strain) for three different vein spacings (*2W* = 0.2, 0.3, 0.4 mm) and for three different times. Upon wounding, as the xylem pressure approaches equilibrium, the neighboring tissue begins to adjust to this new equilibrium by drawing water from their local xylem bundle; this relaxation leads to leaf dilation indicated as percentage changes in colormap. The numerical predictions were simulated for a non-transpiring plant assuming linear elastic response and using experimentally available parameters to evaluate tissue poroelastic diffusivity, *κ*_*t*_ *=*10^−10^ m^2^s^-1^ (See S.I. Section S6). A time-lapse video of the simulation is provided in Movie S4. **c**, Plot of the predicted kinetics of the increase in leaf thickness obtained by averaging the total leaf dilation shown in (b). As the increase in leaf thickness is governed by the poroelastic relaxation timescale of the tissue which depends on the vein spacing (*τ*_*t*_*=W*^*2*^*/κ*_*t*_), the parts of the leaf with denser venation reach equilibrium more quickly. **d**, Comparison of the tissue relaxation timescale obtained from experiments (grey) and the predicted timescale (green) from the model reveals quantitative agreement. Inset: experimental timescale obtained as the mean of the five relaxation timescales from Malone’s experiment shown in (a), with error bars based on the standard deviation; predicted timescale based on a range of values of tissue poroelastic diffusivity based on experimental measurements (Table S2), *κ*_*t*_ = (1±0.5) ×10^−10^ m^2^s^-1^ and a vein spacing of *2W*=0.3 mm, with error bars based on the standard deviation across range of values of *κ*_*t*_.

Following Malone’s observations, we hypothesize that water released at the wound site flowed via the xylem to the tissue of the adjacent leaf in which swelling was observed. Based on this hypothesis, the rapid poroelastic equilibration of the xylem predicted by our model (Fig. 2c) can explain observation (i) of the apparent simultaneous initiation of swelling along the full length of the adjacent leaf. We predict a relaxation time in the xylem, *τ*_*xyl*_ ∼ 40 ms, for wheat with total leaf-to-leaf distance, *L* ∼ 20 cm; this timescale is below the temporal resolution of Malone’s experiment. Proceeding to consider the subsequent stage of the observed swelling, we follow the qualitative suggestion by Malone^8^ that the variation of rates with position along the leaf (observation (ii)) was defined by anatomy: leaves with parallel venation gradually become narrower towards the tip with a commensurate decrease in the spacing between veins. In Fig. 3b, we present the predicted time-evolution of relative dilation in cross-sections of the leaf for three values of vein-spacing (2*W* = 0.2, 0.3, and 0.4 mm) with our poroelastic model of pressure diffusion (Fig. 2b and S.I. Section S6; Movie S4); these spacings correspond to typical values for inter-vein spacing from proximal to distal positions along a wheat leaf.^35^ As seen qualitatively in these colormaps and quantitatively in the integrated relative thickness in Fig. 3c, the predicted evolution of swelling upon initiation from the xylem depends sensitively on the distance, *W* through which pressure changes and fluid must diffuse to relax the tissue. Importantly, we note that our model agrees quantitatively with the timescale of tissue relaxation reported by Malone (*τ*_*t*_ =3.9 ± 1.7 min – Fig. 3d) with no adjustable parameters. This agreement supports the appropriateness of our poroelastic framework for modeling the purely hydraulic dynamics that can be initiated by a wounding event.

### Driving mechanisms for mass flow of chemical elicitors in xylem

Ricca proposed that the primary wounding signal is a chemical elicitor transported with mass flow through the xylem (Fig. 4a).^3^ Although this hypothesis has been supported by several studies, the causation for the underlying mass flow that carries the elicitors remains unclear.^4^ While transpiration is the most obvious potential driving mechanism, mass flow in the xylem of transpiring plants typically ranges from 200 to 400 μm/s,^36,37^ systemic signals can propagate at rates exceeding 1000 μm/s.^25^ Furthermore, wounding signals have been observed to propagate against the direction of the transpiration stream.^38^

Here, we confront predictions of poroelastic mass flows induced by wounding with the observed propagation of signals in calcium reporter plants (*A. th*.; *L* ∼ 1cm; Fig. 4b)^5^ and captured as voltage changes along the leaves of wheat seedlings (*L* ∼ 20 cm, Fig. 4c)^27^. We analyze the distance, *d* [m] that chemical elicitors released at the wound site would be propagated by advection and diffusion in the xylem by: (i) mass flows associated with water being pulled from the xylem into the surrounding tissue during its relaxation (Fig. 2d); and (ii) the steady draw on xylem water by transpiration from the leaf surfaces. As mentioned in discussion of Fig. 2f, the predicted mass flow associated with the relaxation of the xylem itself is too short-lived to displace elicitors over the macroscopic distances that have been observed; we do not model this mode of transmission of chemical elicitors here.

Accounting for the mass flows generated by tissue relaxation (case i) and steady transpiration (case ii), we use our model (see S.I. Section S7) to predict the propagation distance of a chemical elicitor. Assuming that this advected elicitor controls observed local responses along the xylem, we compare this prediction with the experimentally observed propagation of cytosolic calcium (Fig. 4b; see S.I. Section S7.B) and electrical signals (Fig. 4c; see S.I. Section S7.C). Using experimentally available values of tissue and xylem properties and their reported uncertainties (see S.I. Table S2), along with a single fitting parameter for the transpiration rate within a physiological range, the predicted dynamics of signal propagation based on advection-diffusion with mass flows in the xylem are in both qualitative (i.e., shape of saturating displacement curve) and quantitative agreement with experimental measurements. In addition, the model suggests that both water influx at the wound site and transpiration play a role in the transmission of a chemical elicitor, and thus non-transpiring plants should have slower propagation dynamics, as was recently demonstrated experimentally by Gao *et al*.^12^ Counterintuitively, in the wounded leaf, transpiration reinforces mass flow due to poroelastic relaxation of the tissue, with transpiration in the wounded leaf moving toward the stem. This reversal of the direction of transpiration-driven flow in the wounded leaf occurs due to the lower hydraulic resistance of the path along the leaf from the wound relative to that from the soil to the leaf (see S.I. Section S11). Our model predictions provide a strong test of the hypothesis that mass flows resulting from hydromechanical relaxation after wounding can significantly contribute to the propagation of elicitors involved in systemic signaling.

### Propagation of local signals

In contrast to systemic signals, local signals do not involve the xylem bundle as a transmission pathway and can be initiated by wounding a cell that is far from the vasculature as shown in Fig. 5a. Bellandi *et al*.^5^ observed that local calcium signals and apoplastic glutamate (i.e., a potential elicitor) spread radially from the wound site in diffusive-like manner, as depicted as scenario (i) in Fig. 5a and plotted in Fig. 5b.^5^ Moreover, they observed that the dynamics of local calcium signals remained unaffected in plants with plasmodesmal closure achieved by callose deposition, suggesting that calcium signals are most likely triggered by an apoplastic diffusion of an elicitor (e.g., glutamate).

**Fig. 4.**
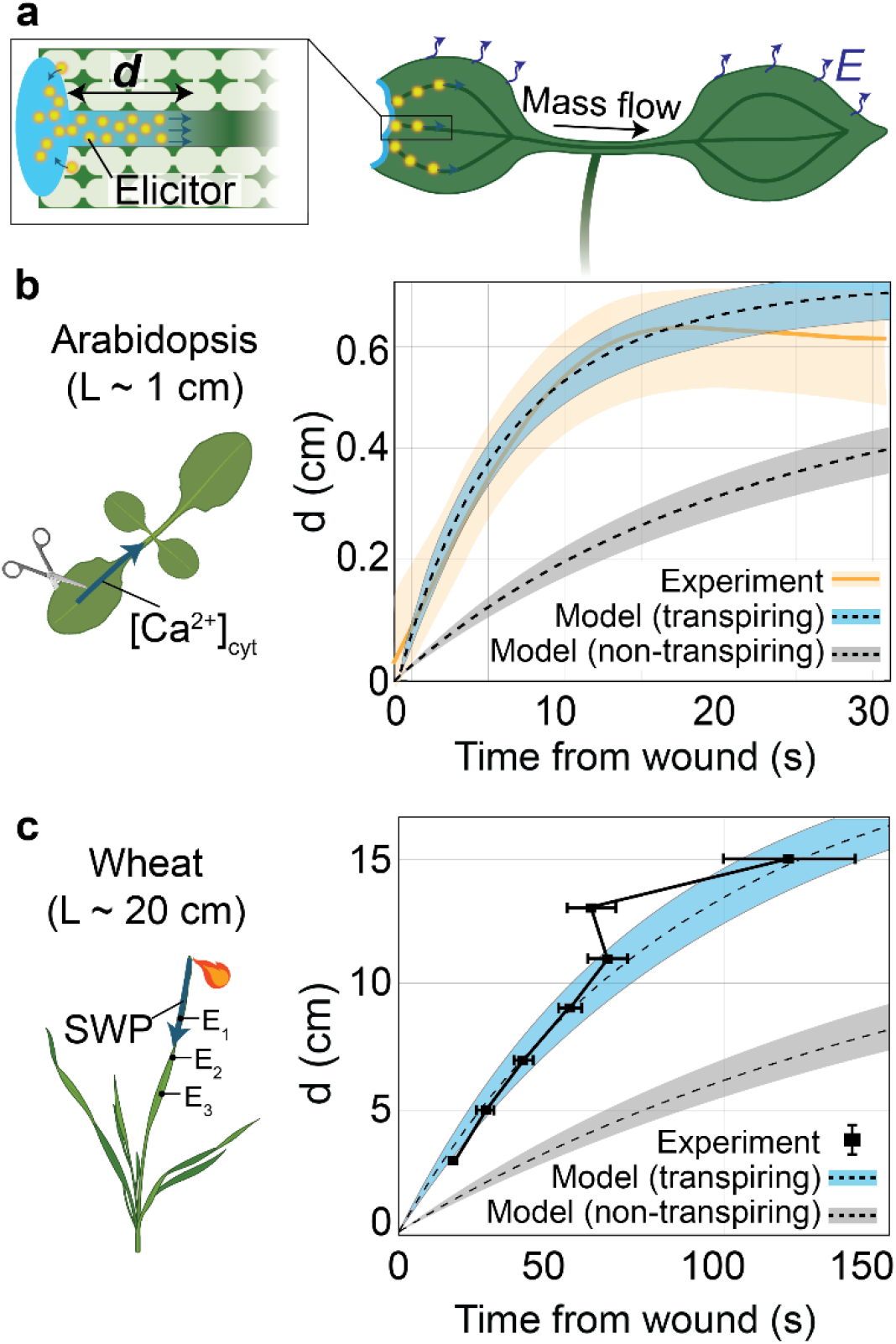
Long-distance advection-diffusion of chemical elicitors with xylem mass flow. **a**, Schematic representation of modeled scenario: mass flow of water released at the wound site from the ruptured cells is driven through the xylem by relaxation of distal tissues (as in Fig. 2d) and by uniform transpiration from the leaf surfaces (*E* [mol m^-2^ s^-1^]). This mass flow in the xylem carries elicitors released at the wound site into non-wounded leaves by advection-diffusion. The propagation distance along the xylem of the elicitors from the wound site, *d* [m] is predicted, as in Fig. 2f. **b**, Experimental measurement from Bellandi *et al*.^5^ of the propagation of wound-induced calcium (Ca^2+^) cytosolic signals along the main xylem vein in an *Arabidopsis thaliana* plant with a genetically-encoded reporter of cytosolic Ca^2+^ and approximate size, *L* ∼ 1 cm. The plot shows experimental measurement of the propagation of calcium signals (in orange), and predicted dynamics for a transpiring (blue shade) plant with transpiration rate of *E*_*0*_ = 1 mmol m^-2^ s^-1^, and non-transpiring plant (grey shade). For the model prediction, we consider a chemical elicitor with diffusivity, *D* = 10^−10^ m^2^s^-1^, and a best fit for the transpiration rate (see S.I. Section S7.B for used parameters). **c**, Experimental measurement from Vodeneev *et al*.^27^ of the propagation of wound-induced electrical signals using three electrodes attached to a wheat seedling with approximate size of *L* ∼ 20 cm. The plot shows experimental measurement of the propagation of electrical signals (black squares), and predicted dynamics of a hypothetical elicitor (blue shaded region) for a transpiring plant with transpiration rate of *E*_*0*_ = 1.2 mmol m^-2^ s^-1^ and for a non-transpiring plant (grey shade). For the model prediction, we consider a chemical elicitor with diffusivity, *D* = 10^−10^ m^2^s^-1^, and a best fit for the transpiration rate (see S.I. Section S7.C for used parameters).

**Fig. 5.**
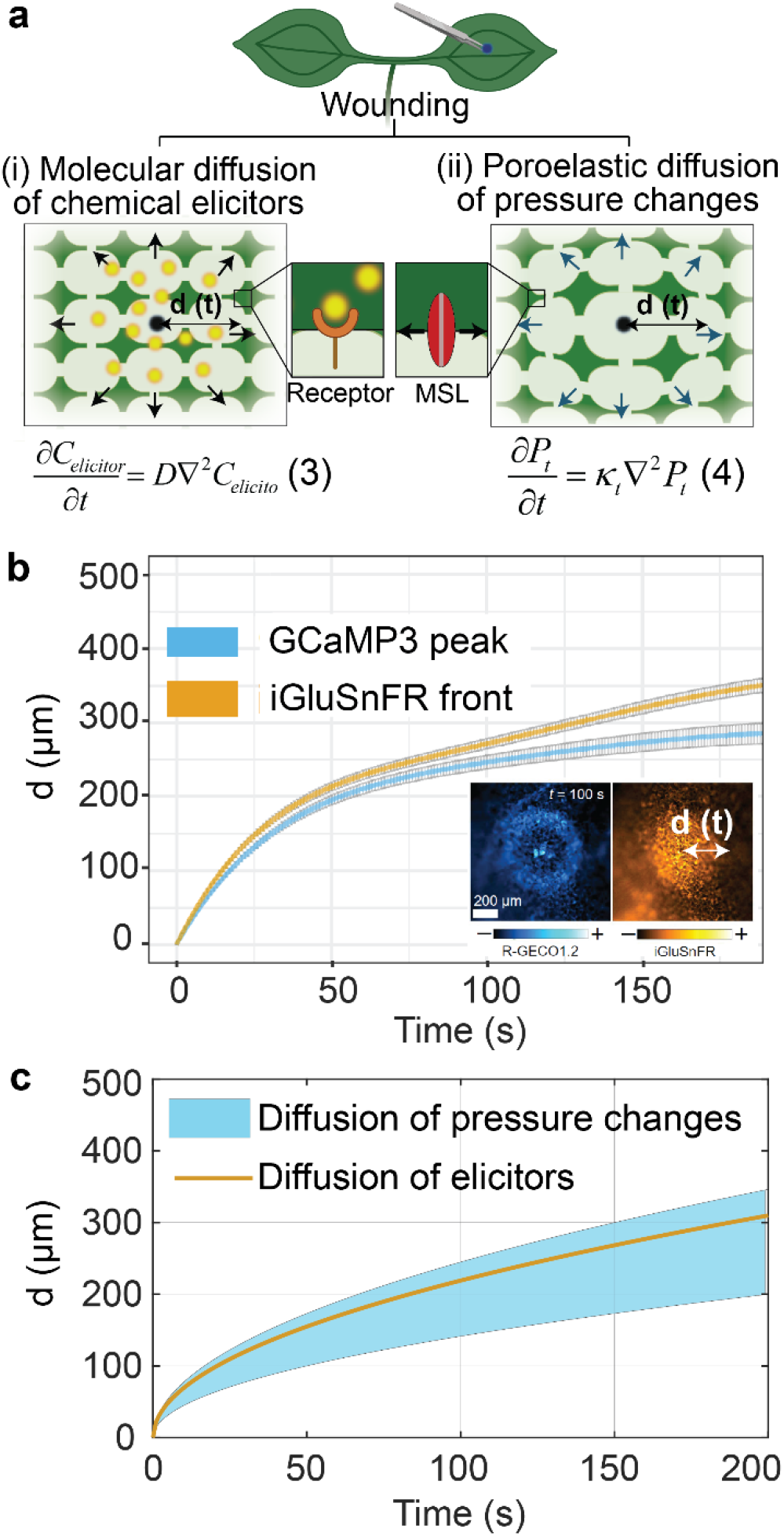
Model of propagation of local signals with diffusion of chemical elicitors or as a direct mechanical response to poroelastic deformation. **a**, Schematic representation of two hypothetical scenarios in which local wounding of the non-vascular tissue occurs far from the vasculature and results in radial propagation (radius, *d* [m]) of: (i) molecular diffusion of elicitors (Eq. 3) with diffusion coefficient of *D* [m^2^s^-1^]) released at wound that can bind to ligand-gated channels or receptors; or (ii) poroelastic diffusion of pressure changes (Eq. 4) with poroelastic diffusivity of *κ*_*t*_ [m^2^s^-1^]), due to release of symplastic water at wound that can induce mechanotransduction of cellular responses, which for example can be sensed by mechanosenstive ion channel (MSL) proteins. **b**, Experimentally observed cytosolic calcium signal and apoplastic glutamate (i.e., potential elicitor) signals observed in independent reporter lines: GCaMP3 (reports cytosolic calcium) and iGluSnFR (reports apoplastic glutamate), where *d* is the radial distance of propagation of the signals (adapted from Bellandi *et al*.^5^). The inset shows fluorescence image for calcium and glutamate response in R-GECO1.2 (reports cytosolic calcium) and iGluSnFR (reports apoplastic glutamate) dual reporter. **c**, Time evolution of the diffusive spread of small chemical elicitor with a diffusion coefficient of *D* =10^−10^ m^2^s^-1^ (Eq. 3, plotted in orange line), and tissue pressure changes (Eq. 4, plotted in blue; see Movie S5) predicted for a range of physiological values of the poroelastic diffusivity (*κ*_*t*_ = (1±0.5) ×10^−10^ m^2^s^-1^) based on independently measured tissue properties (Table S2). The diffusion of a small chemical elicitor released at the wound site, exhibits quantitatively and qualitatively similar behavior to the propagation of pressure changes, thereby creating ambiguity regarding whether the experimentally observed calcium spread is triggered by diffusion of a chemical elicitor released at the wound site, the propagation of pressure changes in the tissue, or both.

Here, we explore alternative or additional potential mechanisms of local propagation associated with poroelastic relaxation initiated by such a wound. First, we exclude the contribution of local advection of chemical elicitors with mass flow in the tissue driven by poroelastic relaxation upon wounding, because the predicted distance of propagation by this mechanism is one order lower than that experimentally observed (see S.I Section S8). Next, we consider direct mechanobiological response to the propagation of pressure changes, as depicted as scenario (ii) in Fig. 5a. Molecular diffusion (Eq. 3 in Fig. 5a, scenario (i)) and poroelastic diffusion of pressure changes (Eq. 4 in Fig. 5a, scenario (ii)) are described by the same form of diffusion equation. Moreover, with experimentally measured values of *k*_*t*_ and *c*_*t*_ (see Table S2), we find that the poroelastic diffusivity of the tissue is similar to molecular diffusivity of small molecules (*D* ≅ *κ*_*t*_ ≅ 10^−10^ m^2^ s^-1^), such that the rate at which changes in pressure propagate in the tissue is similar to that for the diffusion of small molecules.^39^ In Fig. 5c, we demonstrate this similarity of spreading dynamics for molecular diffusion of an elicitor (solid yellow curve) and poroelastic diffusion of changes in pressure (blue region for range of parameters provided in caption). We conclude that the observed dynamics of calcium signals could be explained by the spread of chemical elicitors by molecular diffusion, the spread of pressure changes by poroelastic diffusion direct mechano-transduction, or a combination of both processes. The ambiguity caused by the qualitative and quantitative correspondence between the dynamics of molecular and pressure diffusion in plant tissues presents an important target for additional experiments (see S.I. Section S10).

## Discussion

Hydromechanical phenomena have long been suggested to play a role in plant signaling but the field has lacked a predictive framework to capture these hypotheses with appropriate, physical and physiological considerations.^1,7,40^ Here, we present such a model based on poroelastic responses to wounding and show the predicted dynamics are compatible with observations of wound-induced response (mechanical,^8^ biochemical^5^, and electrophysiological^27^), both qualitatively and quantitatively. This predictive framework suggests that the role of hydraulics on long distance signaling, and therefore other downstream responses, are strongly influenced by global water relations in the plant, including evaporative demand, vascular architecture, and type of wound. This finding highlights the need for their careful consideration of the state of stress and rate of transpiration when comparing signal dynamics across different studies. In particular, the framework can explain the experimentally observed differences between ‘wet wound’ (i.e., immersing the wounded leaf in water) vs ‘dry wound’ (i.e., exposing the wounded leaf to air),^5,41^ by suggesting that dry wounds rapidly deplete the released water, restore xylem tension and reduces elicitor propagation (See S.I. Section S12). The framework also points out to ambiguity between the propagation molecular diffusion and poroelastic diffusion of pressure in non-vascular tissues, calling for further experiments to elucidate the biophysical mechanisms governing local signal transmission, such as simultaneous measurement of local responses (e.g., cytosolic calcium) and tissue pressure (as discussed in S.I. Section S10).

Finally, while our work supports that hydraulics acts as an upstream response, mode of propagation and downstream perturbation for wounding events, for all of these processes, we emphasize that hydromechanical dynamics are generic and non-specific, occurring to some degree in any plant under tension upon local liquid release. This non-specificity suggests that the propagation of hydromechanical perturbation would need to be coupled with other signaling pathways, likely chemical or electrochemical signals, to encode a diverse range of upstream events and downstream responses. This framework lays the groundwork for future models integrating local kinetics of downstream signaling pathways and physiological responses, which could enable predictions regarding the different factors involved in signal initiation, propagation, and target elicitation.

## Methods

### Model inputs

All the model inputs such as hydraulic conductivity (*k*_*x*_, *k*_*t*_), hydraulic capacity (*c*_*x*_, *c*_*t*_), and plant anatomy (*L, W*) are obtained for independent experimental studies on a variety of plant species as discussed in S.I. Section S4 and shown in Table S2. The only fitting parameter we use when comparing experimental results on the propagation of chemical elicitors (Fig. 4) is the transpiration flux, *E*, due to the unknown transpiration status of the observed plants. Nonetheless, the fitted values remain within a physiologically appropriate range (see Table S2).

### Numerical simulation

To numerically solve the governing equations for the xylem (Eq. 1, Fig. 2b) and tissue turgor pressure (Eq. 2, Fig. 2b), we use a finite element analysis software package (COMSOL 6.2, COMSOL Inc., Burlington, MA, USA) as described in S.I. Section S5.

To simulate the predicted increase in leaf thickness shown in Fig. 3b-c, we numerically solve the tissue turgor pressure (Eq. S16) across the leaf thickness using *PDEmodel* solver in MATLAB R2022b (Mathworks, Natick, MA, USA) as described in S.I. Section S6. The MATLAB code of this simulation is provided in Script S1.

To obtain the predicted propagation distance of chemical elicitors released at the wound site (Fig. 4), we first use COMSOL to obtain the resulting xylem and tissue turgor pressure, and then we use MATLAB R2022b to simulate the propagation of chemical elicitors as described in S.I. Section S7. The MATLAB code for these simulations is provided in Scripts S2 and S3, and the COMSOL files are provided in Files F1-F8.

To simulate the propagation of local signals (Fig. 5, Eq. 4), we use the partial differential equations solver *pdepe* in MATLAB R2022b as described in S.I. Section S8. The MATLAB code of this simulation is provided in Script S4.

## Acknowledgements

We thank C. Faulkner and R. Morris for the meticulous review of the manuscript and insightful feedback. We also thank S. Desai, S. Sen and E. Wu for useful discussions. This project was supported by the National Science Foundation STC Center for Research on Programmable Plant System under grant number DBI-2019674, the Air Force Office of Scientific Research under grant number FA9550-18-1-0345, National Science Foundation grant number PHY-2412533, and Agence Nationals de la Recherche under grant number ANR-23-CE30-0046. V. Bacheva was supported by Schmidt Science Fellows, SNF Postdoc.Mobility (grant number 214477), and KIC Postdoctoral Fellowship.

## Notes

### Competing Interest Statement

The authors have declared no competing interest.

