## Supplementary Information for "A unified framework for hydromechanical signaling: Do plant signals go with the flow?"

### This PDF file includes:

### Other supplementary materials for this manuscript include the following:

Movies: Movie S1, Movie S2, Movie S3, Movie S4, Movie S5

MATLAB Scripts: Script S1, Script S2, Script S3, Script S4

COMSOL Files: File F1, File F2, File F3, File F4, File F5, File F6, File F7, File F8

### S1. Existing models on hydromechanical perturbations in plants

In this section, we first provide a brief summary of studies that have modeled hydromechanical perturbations (i.e., involving water flow and transmission of mechanical stresses and strains through plant tissues) in non-wounded plants (Section S1.A). We then discuss studies that focused on modeling the propagation of systemic signals through the xylem following a wounding event, either through the propagation of pressure signals or the propagation of chemical elicitors Section S1.B).

#### A. Poroelastic relaxation of plant tissues in non-wounded plants

To the best of our knowledge, Philip was the first to analyze water flow in plant tissue.[1,2] His initial work focused on modeling water transport through a single cell,[1] which he later extended to the tissue scale by considering a linear collection of cells.[2] At the tissue scale, Philip modeled water flow through cells by considering cell-to-cell permeability. He assumed that changes in cell volume are elastic and small, and that there is a linear relationship between changes in cell volume and water potential. Philip demonstrated that water flow can be mathematically described as a diffusion phenomenon, with the diffusion coefficient dependent on cell permeability and the elastic properties of the cell wall—a coefficient now referred to as the poroelastic diffusion coefficient. Rockwell *et al.*[3] revisited Philipp's work, incorporating additional apoplastic pathways (via the cell wall) alongside cell-to-cell pathways. They derived coupled governing equations in which the water potential in the symplast and apoplast are coupled but distinct (not in local equilibrium). They also developed a composite model with the assumption of local equilibrium of water potential between symplast and apoplast (as in Eq. 2 in Fig. 2b of main text – see Section S2.C and S3.A for details) and found that it agreed with the two-compartment model with ~10% in the prediction of relaxation of water potential in a macroscopic tissue (e.g., leaf rehydration). In this work, we adapt and extend the poroelastic models of Philip and Rockwell in developing our poroelastic framework for the treatment of scenarios relevant to long-distance signaling.

Louf *et al.*[4] investigated how pressure changes propagate in a plant branch when it is bent. They developed both a model and experiments using plant branches as well as a synthetic poroelastic branch made from a PDMS cylinder perforated with longitudinal channels. Their simple energetic model showed that when bending occurs, it is energetically favorable for the poroelastic branch to generate transverse strain, squeezing its cross-section and resulting in a reduction in volume. Since the branches are filled with water (an incompressible fluid), this volume change is converted into an overpressure proportional to the bulk modulus of the branch and the tube diameter. While this work demonstrates that plant branches behave as poroelastic materials, it does not account for water transfer from the vasculature to the surrounding tissue and focuses only on unwounded branches.

**Differences between existing models and our framework.** Compared to the existing models of poroelastic relaxation in non-wounded plants presented above, our framework provides a unified approach that:

1. Considers the case of a wounded plant and provides the boundary conditions resulting from a wounding event.
2. Solves for the poroelastic dynamics in two separate compartments composed of xylem bundles and non-vascular tissue.
3. Considers explicit representation of venation and tissue architecture over whole-plant scales to predict spatial dynamics of pressure, flow, and strain during transient of xylem and tissue relaxation.
4. Considers the impact of non-negligible solute concentration in the apoplast (Section S3.A and S3.B) to account for the specific scenario of wounding in which solutes can enter the apoplast from the disrupted symplast or the exogenous application to the wound site.
5. Accounts for uniform and constant transpiration from the tissue.

### B. Propagation of systemic signals in wounded plants

**Propagation of hypothetical pressure signals.** Evans and Morris[5] treated the xylem as a single isolated cylindrical vessel with elastic walls containing an incompressible and non-viscous fluid. We note that their assumption of a non-viscous fluid with an inertially dominated response places their model in the opposite limit with respect to viscous and inertial responses than is appropriate for water in conduits of the radial dimension typical of xylem at the expected flow speeds. More precisely, the non-dimensional ratio of inertial to viscous forces is captured by the Reynolds number: $Re = ud/\nu$, where in xylem, the signal velocity[6], $u = 10^{-3} - 10^{-2}[m\,s^{-1}]$, the xylem diameter, $d = 10^{-5} - 10^{-4}$[m], and the kinematic viscosity of water, $\nu = 10^{-6}[m^2\,s^{-1}]$, such that $Re = 10^{-2} - 1$. In this limit of low Reynolds number, the viscous response of water in the xylem is expected to be completely dominant.[7] With their model of inertially dominated relaxation, Evans and Morris predicted that the velocity of propagation of pressure changes in the xylem is above 10 m $s^{-1}$, which, as they note, is much higher than the experimentally observed velocity of electrical signals that is on the order of 1-10 mm $s^{-1}$. Based on this mismatch in velocities, they suggested that propagation of pressure changes is unlikely to be the trigger for the propagation of electrical signals. In this study, we develop a model (see Section S2) for the physically expected regime in which viscous and elastic stresses dominate (i.e., a poroelastic model).

Sukhova *et al.*[8] proposed a simplified model of the transient of pressure change in the xylem after wounding. Their approach neglected the elastic nature of the xylem tissue by considering a rigid tube and avoided explicitly considering the time-dependent momentum balance. To capture dynamics without explicitly modeling it, they introduced a relaxation time as a fitting parameter over which they allowed a steady state, linear profile of pressure along the tube to decay. This approach does not provide a basis for predicting (i.e., based on material properties, geometry, and fundamental conservation and rate laws) either the characteristic relaxation time or the spatial evolution of the pressure field. As such they simply assumed that the relaxation time was on the order of tens of seconds. As we have discussed in the main text, this assumption does not match the prediction of a formal treatment of the expected poroelastic response of the xylem (~10 ms for a 1 m long plant, see Fig. 2e).

**Propagation of hypothetical chemical elicitors with mass flow.** Two studies have focused on modeling the advective processes that govern the propagation of chemical elicitors in the xylem. Blyth and Morris[9] proposed that both advective and diffusive processes in single xylem vessels may account for experimentally observed propagation of electrical and calcium signals. They used Taylor-Aris dispersion to show that predictions from the model are in good agreement with experimentally observed electrical dynamics. While this work provides a solid foundation of the expected advection processes in xylem flow following wounding events, as the authors acknowledge, their model lacks an explanation for the causation for the underlying mass flow.

Vodeneev *et al.*[10] suggested that the driving mechanism of a chemical elicitor in the xylem involves "turbulent" diffusion arising from rapid pressure changes that enhance the dispersal of chemical elicitors. This proposal was motivated by their observation that propagation speeds were much larger than expected for molecular diffusion of a small elicitor. However, it is unlikely that turbulence could occur in typical low-Reynolds conditions (i.e., where turbulence cannot develop) of the xylem.[11]

**Differences between existing models and our framework.** As discussed in this section, existing works have focused on either the modeling of mechanical (i.e., propagation of pressure changes) or advective processes (i.e., propagation of elicitors) in systemic signals. Additionally, these models have neither addressed the coupling of the xylem with non-vascular tissues nor investigated the mechanisms of the underlying mass flow. Our framework provides a unified approach that:

1. Considers an explicit representation of the coupling of xylem and tissue architectures on the whole-plant scale, including root, stem, and leaves.
2. Integrates both advective and mechanical processes following a wounding event.
3. Derives the expected propagation dynamics of systemic and local signals by solving governing equations for spatially and temporally resolved predictions of pressure, mass flow, and advection of elicitors.
4. Offers a plausible explanation of the mass flow that can advect chemical elicitors away from the wound site.

5. Considers the impact of non-negligible solute concentration in the apoplast to account for the specific scenario of wounding in which solutes can enter the apoplast from the disrupted symplast or the exogenous application to the wound site.
6. Defines all model inputs via physiologically relevant and experimentally measured parameters such as hydraulic conductivity and capacitance, plant anatomy, and transpiration status.

### S2. Derivation of poroelastic framework of plant signaling

In this section, we provide the derivation of the poroelastic framework. We begin by discussing the main assumptions that enable us to make predictions on the experimentally available data shown in the main text (Section S2. A). We then derive the poroelastic equations that govern the propagation of systemic (Section S2. B) and local (Section S2. C) pressure signals, and we discuss the behavior of these equations through scaling analysis. We conclude by discussing how the framework can be generalized to cases beyond the ones discussed in this work (Section S3).

#### A. Framework statement and assumptions

To derive the poroelastic framework to make predictions on the experimentally available observations on plant signaling, discussed in the main text, we make the following assumptions (Fig. S1):

- Assumption 1: Treat leaves as thin, plate-like structures, with leaf thickness much smaller than their width and length, composed of two compartments: xylem bundles and tissue.
- Assumption 2: Neglect variation in the osmotic pressure of the tissue due to actively regulated processes and changes in cell volume.
- Assumption 3: Treat the xylem bundles as formed of a collection of xylem vessels that are modeled as hollow tubes.
- Assumption 4: Treat the tissue as 'composite' material in which the symplast and apoplast are in local equilibrium.
- Assumption 5: Treat the xylem bundle and tissue as connected via a semi-permeable membrane that is perfectly impermeable to solutes.
- Assumption 6: Treat the wounding event as resulting in the release of an excess of available liquid water with a negligible concentration of solutes.

In Section S3, we discuss how the framework can be generalized beyond assumptions 2, 4, 5 and 6.

##### Model geometry and water potential

Fig. S1 illustrates the model under consideration focusing on two leaves that share a direct vascular connection. We model the leaves as a 2D plate with uniform leaf thickness, $d_{th}$ [m], comprising a xylem bundle (length $L$ [m] and width $l_{xyl}$ [m]) sandwiched between non-vascular tissue (length $L$ and vein spacing $2W$ [m]). The xylem bundle, which is composed of a collection of individual xylem vessels

**Table S1.** Symbol definition.

| Quantity | Symbol | Units |
|---|---|---|
| Water content | $M_w$ | mol $m^{-3}$ |
| Solute concentration | $C$ | mol $m^{-3}$ |
| Water potential | $\psi$ | Pa |
| Transpiration flux | $E$ | mol $m^{-2}$ $s^{-1}$ |
| Turgor pressure | $P$ | Pa |
| Osmotic pressure | $\pi$ | Pa |
| Water flux | $J$ | mol $m^{-2}$ $s^{-1}$ |
| Hydraulic conductivity | $k$ | mol $m^{-1}$ $Pa^{-1}$ $s^{-1}$ |
| Xylem-tissue permeability | $L_{x\text{-}t}$ | m $Pa^{-1}$ $s^{-1}$ |
| Root-stem transport coefficient | $h_{stem}$ | mol $m^{-2}$ $Pa^{-1}$ $s^{-1}$ |
| Hydraulic capacity | $c$ | mol $m^{-3}$ $Pa^{-1}$ |
| Poroelastic diffusivity | $\kappa$ | $m^2$ $s^{-1}$ |
| Molar volume of water | $\bar{v} = 1.8\times10^{-5}$ | $m^3$ $mol^{-1}$ |
| Dynamic water viscosity at 25C | $\eta = 8.9\cdot10^{-4}$ | Pa s |

with radius, $l_{ves}$, and adjacent cells from the tissue are treated as linear and isotropic poroelastic media. The leaf is transpiring from the top and bottom side of the tissue with a constant transpiration flux, $E$ [mol $m^{-2}$ $s^{-1}$]. We represent the node where the two leaves are connected to the stem as a water source that replenishes the water lost through transpiration.

The water flow in poroelastic media depends on the water potential, $\psi$ [MPa], which characterizes the availability of water for chemical and physical transformation. Neglecting the gravitational contribution, we identify two contributions to the water potential of the xylem, $\psi_{xyl}$ and the tissue, $\psi_t$:

$$\psi_{xyl} = P_{xyl} - \pi_{xyl} = P_{xyl} - RTC_{xyl}, \tag{S1}$$

$$\psi_t = P_t - \pi_t = P_t - RTC_t, \tag{S2}$$

where $P_{xyl}$ and $P_t$ [Pa] are the xylem and tissue turgor pressure, respectively relative to the atmospheric pressure, $R$ [J $mol^{-1}$ $K^{-1}$] is the ideal gas constant, $T$ [K] is the temperature, and $C_{xyl}$ and $C_t$ [mol $m^{-3}$] are the concentration of solute in the xylem and tissue, respectively. The second term, $\pi$ [Pa] is known as the osmotic pressure and is given by the van't Hoff relation, $\pi=RTC$.[12] The xylem osmotic pressure in a non-wounded plant is typically negligible due to its low osmotic content ($\pi_{xyl}$ = 0 MPa).

**Assumption 1.** Plant leaves typically exhibit thin, plate-like structures with leaf thickness much smaller than the separation between vessels: $d_{th} << W$ and $L$. Given this geometry, we assume that water potential through its thickness ($z$-axis) remains uniform in both the xylem bundle and surrounding tissue. This assumption is justified based on the scaling of the transients of gradients of pressure predicted in a poroelastic material with the distance of propagation (Eqs. S27-28): $\frac{\tau_{d_{th}}}{\tau_W} = \frac{d_{th}^2}{W^2}$ and $\frac{d_{th}^2}{L^2} \ll 1$, such that gradients through the thickness will decay much more rapidly than those across the lateral dimensions of the leaves.

**Assumption 2.** We assume that the osmotic pressure in the tissue prior to wounding is spatially uniform across all cells. Following a wounding event, we assume that during the propagation of relatively fast (seconds to minutes) wound-induced responses, there are no actively regulated processes that adjust the solute concentration in the tissue (See Section S3.C for the discussion on how the framework can be adjusted to consider actively regulated processes that adjust the osmotic pressure in the tissue). Therefore, the number of solutes in each cell within our composite model of the tissue remains fixed. Moreover, we consider only small deformations of the cells in the x-y plane as they take up water following a wounding event, and therefore assume that the cell volume remains nearly constant. This assumption is supported by experimental observations of tissue swelling following wounding, as observed by Malone,[13] in which the total increase in leaf thickness is on the order of 10 µm; this uniaxial and volumetric deformation is less ~5% for a total leaf thickness of ≅ 200 µm. Based on these two assumptions (fixed solute number and negligible cell dilation effects), the osmotic pressure in the symplast and apoplast of the tissue remains constant ($\pi_t = RTC_t$ = const), such that changes in tissue water potential are reflected as changes in turgor pressure, $P_t$.

**Assumption 3.** We assume that xylem vessels are hollow circular tubes, in which the flow follows Poiseuille's law. As a result, the hydraulic conductivity in each vessel, $k_{ves}$ [mol m$^{-1}$ Pa$^{-1}$ s$^{-1}$] is:

$$k_{ves} = \frac{l_{ves}^2}{8\bar{v}\eta}, \tag{S3}$$

where $\bar{v}$ [m$^3$ mol$^{-1}$] is the molar volume of water, and $\eta$ [Pa s] is the dynamic water viscosity.

The water flux through the xylem bundles, $\boldsymbol{J}_{xyl}$ [mol m$^{-2}$ s$^{-1}$] occurs along gradients of xylem pressure and follows Darcy's law:

$$\boldsymbol{J}_{xyl}(x,y,t) = -k_{xyl}\left(\frac{\partial P_{xyl}(x,y,t)}{\partial x}\hat{\boldsymbol{x}} + \frac{\partial P_{xyl}(x,y,t)}{\partial y}\hat{\boldsymbol{y}}\right) \tag{S4}$$

where $k_{xyl}$ [mol m$^{-1}$ Pa$^{-1}$ s$^{-1}$] is the hydraulic conductivity of the xylem bundle. Assuming that the xylem bundle is an isotropic medium with porosity, ε, (i.e., the ratio between vessel volume and the total xylem bundle volume), the hydraulic conductivity of the xylem can be defined through the vessel conductivity as[14]:

$$k_{xyl} = k_{ves}\frac{\varepsilon}{3} = \frac{\varepsilon l_{ves}^2}{24\bar{v}\eta}. \tag{S5}$$

**Assumption 4.** We treat the tissue as a 'composite' material or effective medium in which the symplast and apoplast are in local equilibrium ($\psi_{sym} = \psi_{apo}$). This assumption simplifies the analysis with only a minor trade-off in accuracy. We estimate an error of less than 5% compared to treating the symplast and apoplast as separate domains, as discussed in Section S3.A. We assume that the water in the cells of the tissue occurs via symplastic (plasmodesmata-mediated) and cross-membrane (plasma membrane-mediated, e.g., aquaporin-mediated) pathways. Following previous studies,[3,15] we neglect the contributions of the apoplastic pathways (i.e., flow through the cell-wall) as relatively small areas are available for apoplastic flows such that the conductance of this path is small relative to those in the symplast. The water through the symplastic pathways occurs along gradients of pressure, and the water through the cross-membrane occurs through gradient of water potentials due to the presence of semi-permeable membranes. However, based on our assumption that osmotic pressure in the tissue remains constant and uniform (assumption 2), the water flux through the tissue, $\boldsymbol{J}_t$ [mol m$^{-2}$ s$^{-1}$] occurs only along gradients of turgor pressure:

$$\boldsymbol{J}_t(x,y,t) = -k_t\left(\frac{\partial P_t(x,y,t)}{\partial x}\hat{\boldsymbol{x}} + \frac{\partial P_t(x,y,t)}{\partial y}\hat{\boldsymbol{y}}\right), \tag{S6}$$

where $k_t$ is the hydraulic conductivity of the tissue that includes contribution from the two parallel pathways: symplastic, $k_{t,PD}$ and cross-membrane, $k_{t,PM}$:

$$k_t = k_{t,PD} + k_{t,PM}. \tag{S7}$$

**Assumption 5.** We assume that for liquid water (sap) to transfer between the xylem (apoplast) and the tissue compartment it must pass into the symplastic space of the tissue via semi-permeable plasma membranes that are perfectly impermeable to solutes. This assumption is consistent with our assumption that transport in the tissue is dominated by the symplastic and transmembrane paths (assumption 4). Therefore, the flux of water across these membranes depends on the difference of the water potential between the xylem and tissue:

$$J_{x-t}(x,t) = \frac{L_{x-t}}{\bar{v}}\left(\psi_{xyl}(x,y,t) - \psi_t(x,y,t)\right)\Big|_{xylem-tissue}, \tag{S8}$$

where $L_{x-t}$ [m Pa$^{-1}$ s$^{-1}$] is the permeability between the xylem bundle and the adjacent cells in the tissue.

**Assumption 6.** We assume that the wounding event results in the release of a negligible concentration of solutes with respect to affecting the water potential in the xylem (i.e., $\pi_{\mathrm{xyl}} \cong 0$ MPa) and an excess of available liquid water at the wound site:

$$P_{xyl}(x,y,t>0)\Big|_{wound} = P_t(x,y,t>0)\Big|_{wound} - \pi_t = 0. \tag{S9}$$

See Section S3. B for the case where at the wound site the concentration of solutes is non-negligible.

### B. Systemic signals

We define systemic signals as the signals that propagate via the xylem to the rest of the plant tissue following the wounding of the tip of one of the leaves by cutting, burning, or squeezing the vasculature. Following Rockwell *et al.*[3], we treat xylem bundles and tissue as a poroelastic media composed of a solid elastic matrix filled with liquid. We assume that during water uptake, we are in the limit of small strains of the matrix and thus we neglect displacement of the matrix. Consistent with Rockwell's notations, we track the water content of a reference volume of poroelastic media (either xylem or tissue), expressed as moles of water per unit volume, denoted as $M_w$ [mol m$^{-3}$]. While poroelastic models often use mass fraction (i.e., mass of water per total mass of the poroelastic media) to track changes in water content, we have chosen to use moles of water per unit volume for consistency with Rockwell's work and the experimentally reported values on tissue properties.

Mass conservation requires that changes in the concentration of water molecules, $M_w$ [mol m$^{-3}$] in the xylem bundle and tissue, are equal to net flux of water, or:

$$\frac{\partial M_{w,xyl}(x,y,t)}{\partial t} = -\nabla \cdot \boldsymbol{J}_{xyl}(x,y,t), \tag{S10}$$

$$\frac{\partial M_{w,t}(x,y,t)}{\partial t} = -\nabla \cdot \boldsymbol{J}_t(x,y,t) - \frac{2E}{d_{th}}. \tag{S11}$$

In Eq. S11, we include uniform transpiration, $E$ [mol m$^{-2}$ s$^{-1}$] from the surfaces of the leaves as a constant sink for water from the tissue.

Due to mass conservation, the fluxes at the xylem-tissue interface are equal:

$$-k_{xyl}\frac{\partial P_{xyl}}{\partial y}(x,y,t)\Bigg|_{xylem-tissue} = -k_t\frac{\partial P_t}{\partial y}(x,y,t)\Bigg|_{xylem-tissue} = J_{x-t}(x,t). \tag{S12}$$

Following the work of Rockwell *et al.*,[3], and based on the assumption of negligible changes in osmotic pressure, we find the relation between pressure and water content as:

$$\frac{\partial M_{w,xyl}}{\partial P_{xyl}} = c_{xyl}, \tag{S13}$$

$$\frac{\partial M_{w,t}}{\partial P_t} = c_t, \tag{S14}$$

where $c_{xyl}$ and $c_t$ [mol m$^{-3}$ Pa$^{-1}$] are the hydraulic capacities of the xylem bundles and the tissue related to the elastic properties of the cell wall. Assuming that the hydraulic conductivities remain constant across the leaf, the conservation equations for water (Eqs. S10 and S11) with the flux laws (Eqs. S4 and S6) give the following governing equations for the xylem pressure ($P_{xyl}$) and turgor pressure ($P_t$):

$$\text{Xylem:} \frac{\partial P_{xyl}(x,y,t)}{\partial t} = \kappa_{xyl}\left(\frac{\partial^2 P_{xyl}(x,y,t)}{\partial x^2} + \frac{\partial^2 P_{xyl}(x,y,t)}{\partial y^2}\right) = \kappa_{xyl}\nabla^2 P_{xyl}(x,y,t), \tag{S15}$$

$$\text{Tissue:} \frac{\partial P_t(x,y,t)}{\partial t} = \kappa_t\left(\frac{\partial^2 P_t(x,y,t)}{\partial x^2} + \frac{\partial^2 P_t(x,y,t)}{\partial y^2}\right) - \frac{2E}{c_t d_{th}} = \kappa_t \nabla^2 P_t(x,y,t) - \frac{2E}{c_t d_{th}}. \tag{S16}$$

In Eqs. S15 and S16, $\kappa_{xyl}$ and $\kappa_t$ [m$^2$s$^{-1}$] are the poroelastic diffusivities for axial propagation of changes in pressure along the xylem and lateral propagation of changes in pressure in the tissue. These definitive parameters for poroelastic dynamics are given by the ratio of the hydraulic permeability and capacitance of the respective tissue:

$$\kappa_{xyl} = \frac{k_{xyl}}{c_{xyl}}, \tag{S17}$$

$$\kappa_t = \frac{k_t}{c_t}, \tag{S18}$$

Eqs. S15 and S16 have the form of diffusion equations with poroelastic diffusivities, $\kappa$ [m$^2$s$^{-1}$] that are analogous to a heat equation, with the poroelastic diffusivities analogous to a thermal diffusivity in that it arises from the ratio of a conductivity to a local storage capacity.

To complete the model definition, we need to provide the boundary and initial conditions. We assume that the wounding event results in the release of a negligible concentration of solutes and an excess of available liquid water at the wound site (assumption 6, Eq. S9):

$$P_x(x,y,t>0)\big|_{wound} = P_t(x,y,t>0)\big|_{wound} - \pi_t = 0. \tag{S19}$$

See Section S3.B for the case where at the wound site the concentration of solutes is non-negligible.

For the other boundaries defined by the leaf margin (see Fig. S1b), we assume no-flux conditions:

$$\nabla P_t(x,y,t)\cdot \hat{\boldsymbol{n}} = 0. \tag{S20}$$

where $\hat{\boldsymbol{n}}$ is the normal vector at the leaf margin. We model the node at which the leaf petioles are connected to the stem (see Fig. S1a-b) as a square domain ($l_{xyl}$ x $l_{xyl}$) in the xylem bundle, and we impose flux boundary condition of water with a variable flux on all boundaries of the node domain:

$$-k_{xyl}\left.\frac{\partial P_{xyl}(x,y,t)}{\partial x}\right|_{node} = h_{stem}\cdot\left(\psi_{soil} - P_{xyl}(x=0,y=0,t>0)\right), \tag{S21}$$

where $h_{stem}$ [mol m$^{-2}$ Pa$^{-1}$ s$^{-1}$] is a transport coefficient that represents the total conductance of the stem and roots, and $\psi_{soil}$ is the water potential in the soil. Assuming a well-watered plant ($\psi_{soil}$ = 0), this boundary condition can be simplified:

$$k_{xyl}\left.\frac{\partial P_{xyl}(x,y,t)}{\partial x}\right|_{node} = h_{stem}\cdot P_{xyl}(x=0,y=0,t>0). \tag{S22}$$

While the initial condition depends on whether the plant is transpiring, it is important to note that the timescales of the transient response do not depend on the initial condition. Instead, these timescales are determined by the properties of the xylem and tissue, and the plant size, as derived in the *Scaling Analysis* below.

**Non-transpiring plant**. For a non-transpiring plant, we assume that the water potential in the xylem bundles and tissue is at equilibrium and constant:

$$P_{xyl}(x,y,t<0) = P_t(x,y,t<0) - \pi_t = P_0 = \text{const.} \tag{S23}$$

**Transpiring plant**. To find the initial xylem and tissue pressure, we solve numerically the steady state of the governing equations, Eqs. S15 and S16, with non-flux boundary condition at all leaf margins including the wound site using COMSOL (See Section S5 for details on the COMSOL simulation). We assume that water potential initially in the whole plant is in equilibrium and constant (Eq. S23). Fig. S2 shows a simulation plot of the resulting initial xylem and tissue turgor pressure. As expected, the xylem pressure drops from the stem toward the tip of both leaves, giving rise to a pressure gradient that drives the transpiration flux. This initial condition is used in the simulations of dynamics presented in Fig. 2.

**Scaling analysis.** In order to assess the importance of individual terms of the governing equations, we non-dimensionalize them using the following change of variables:

$$\bar{P} = \frac{P}{P_0}, \bar{x} = \frac{x}{L}, \bar{y} = \frac{y}{W}, \bar{t} = \frac{t}{\tau}. \tag{S24}$$

Due to the narrow anatomy of xylem bundles, the typical width of xylem bundles is much smaller than the length of a bundle ($l_{xyl} \ll L$), and thus the variations in xylem water potential across the y-direction are negligible. Substituting the variables into the governing equations, Eqs. S15 and S16, we get:

$$\text{Xylem:} \frac{\partial \bar{P}_{xyl}\left(\bar{x},\bar{y},\bar{t}\right)}{\partial \bar{t}} = \frac{\tau}{L^2/\kappa_{xyl}} \frac{\partial^2 \bar{P}_{xyl}\left(\bar{x},\bar{y},\bar{t}\right)}{\partial \bar{x}^2}, \tag{S25}$$

$$\text{Tissue:} \frac{\partial \bar{P}_{t}\left(\bar{x},\bar{y},\bar{t}\right)}{\partial \bar{t}} = \frac{\tau}{W^2/\kappa_t} \left( \frac{W^2}{L^2} \frac{\partial^2 \bar{P}_t\left(\bar{x},\bar{y},\bar{t}\right)}{\partial \bar{x}^2} + \frac{\partial^2 \bar{P}_t\left(\bar{x},\bar{y},\bar{t}\right)}{\partial \bar{y}^2} \right) - \frac{2E\tau}{P_0 c_t d_{th}}. \tag{S26}$$

From this scaling analysis, we identify that propagation of changes in the xylem pressure is characterized by a poroelastic diffusive timescale, $\tau_t$ [s] defined as:

$$\tau_{xyl} = \frac{L^2}{\kappa_{xyl}}, \tag{S27}$$

In the case of long-distance signal propagation ($L \gg W$), the propagation of changes in the tissue turgor pressure is characterized by a poroelastic diffusive timescale defined by $\tau_t$:

$$\tau_t = \frac{W^2}{\kappa_t}. \tag{S28}$$

#### C. Local signals

In contrast to systematic signals, local signals do not involve the xylem bundle as a transmission pathway and can be initiated by wounding a cell that is far from the vasculature as shown in Fig. S3. To model the propagation of local pressure signals in the tissue following the wounding of a central cell with radius, $r_{wound}$ [m], we here assume that all cells constitute an effective medium that treats the tissue as a single fluid-carrying material with uniform properties (a 'composite' model). In Section S3.A, we treat both symplast and apoplast separately, and we show that the 'composite' model is a good estimate of the full set of coupled equations of the symplast and apoplast.

Based on the assumption of negligible changes in the tissue osmotic pressure, the governing equation of the pressure in the tissue in cylindrical coordinates for the composite model is:

$$\frac{\partial P_t(r,t)}{\partial t} = \kappa_t \left( \frac{1}{r} \frac{\partial P_t(r,t)}{\partial r} + \frac{\partial^2 P_t(r,t)}{\partial r^2} \right). \tag{S29}$$

Importantly, we note that this equation has the identical form as the equation for molecular diffusion equation of solutes released at the wounded cells, with diffusivity, $\kappa_t$. Prior the wounding event, we assume that the water potential in the tissue is at equilibrium and constant:

$$P_t(r,t<0)-\pi_t = P_0. \tag{S30}$$

We assume that the wounding event results in a release of available liquid water at the wounded cell with negligible concentration of solutes:

$$P_t(r = r_{wound}, t>0)-\pi_t = 0. \tag{S31}$$

See Section S3.A for the effect of solutes released at the wound site. Far from the wound ($r \gg r_{wound}$), we assume no-flux condition:

$$\frac{\partial P_t}{\partial r}\left(r \gg r_{wound}, t>0\right) = 0. \tag{S32}$$

We solve Eqs. S29-32 numerically to generate the predictions presented in the main text (Fig. 5 and Movie S5), as described in detail in Sections S8.

**Scaling analysis.** In order to estimate the propagation of the pressure changes in the tissue as function of time, we non-dimensionalize the governing equation Eq. S29 using the following change of variables:

$$\bar{P}_t = \frac{P_t}{P_0}, \bar{r} = \frac{r}{d}, \bar{t} = \frac{t}{\tau}, \tag{S33}$$

where $d$ [m] is the radius of the radial propagation of the water potential at time $t = \tau$. Substituting the variables into the governing equations Eq. S29 we get:

$$\frac{\partial \bar{P}_t(\bar{r},\bar{t})}{\partial \bar{t}} = \frac{\tau}{d^2/\kappa_t}\left(\frac{1}{\bar{r}}\frac{\partial \bar{P}_t(\bar{r},\bar{t})}{\partial \bar{r}} + \frac{\partial^2 \bar{P}_t(\bar{r},\bar{t})}{\partial \bar{r}^2}\right). \tag{S34}$$

From the scaling analysis, we obtain that radial propagation of the water potential in the tissue depends on the square root of time, $d = \sqrt{\kappa_t t}$ .

### S3. Generalization of framework

#### A. Effect of solutes on local signals

In this section, we derive the coupled transport equations in the tissue with explicit, coupled apoplastic and symplastic compartments. It is important to note that here, 'apoplast' refers to the extracellular space outside of plant cell membranes, and not the xylem, which is also often referred to as part of the apoplast. This case allows us to treat scenarios in which the assumption of local equilibrium of water potential made in our composite model (Section S2.C, assumption 4, Eq. S29) does not hold. We use this generalized case to consider the scenario in which a wounding event results in a release of water in the apoplast with a non-negligible concentration of solutes. We then compare the prediction of the water potential dynamics with a composite model, where we assume that the wounding results in the release of water with negligible solute concentration of solutes.

Due to the symmetry of the local wounding, we here consider the one-dimensional transport in the radial direction to capture the spread of the water potential, as shown in Fig. S4. We assume that the tissue cells are composed of symplast with a thickness, $l_{sym}$ [m] and width, $l_{sym}$, and apoplast (i.e., cell wall) with a thickness $l_{apo}$, width, $l_{sym}$. The water potential in the apoplast, $\psi_{apo}$ and symplast, $\psi_{sym}$ are given by:

$$\psi_{apo} = P_{apo} - RTC_{apo} = P_{apo} - \pi_{apo}, \tag{S35}$$

$$\psi_{sym} = P_{sym} - RTC_{sym} = P_{sym} - \pi_{sym}, \tag{S36}$$

where $P_{apo}$ and $P_{sym}$ [Pa] are the apoplastic and symplastic pressures, respectively, and $C_{apo}$ and $C_{sym}$ [mol m$^{-3}$] are the concentration of the solutes in the apoplast and symplast, respectively. As discussed in Section S2.A (assumption 2), we assume that the osmotic pressure in the symplast remains constant, $RTC_{sym}$ = const. Relative to Eq. S19, in Eqs. S35 and S36, we allow for non-negligible osmotic potential in the apoplast and we do not assume equilibrium between the apoplast and symplast.

Following the wounding event, the released symplastic water can propagate as an apoplastic flux, $J_{apo}$ [mol m$^{-2}$ s$^{-1}$], a symplast flux, $J_{sym}$, and a flux between the apoplast and the symplast, $J_{apo\text{-}sym}$:

$$\boldsymbol{J}_{\boldsymbol{apo}}(r,t) = -k_{apo}\frac{\partial P_{apo}(r,t)}{\partial r}\hat{\boldsymbol{r}}, \tag{S37}$$

$$\boldsymbol{J}_{\boldsymbol{sym}}(r,t) = -k_{sym}\frac{\partial P_{sym}(r,t)}{\partial r}\hat{\boldsymbol{r}}, \tag{S38}$$

$$\begin{aligned} J_{apo-sym}(r,t) &= \frac{L_{apo-sym}}{\bar{v}}\left(\psi_{apo}(r,t) - \psi_{sym}(r,t)\right) = \\ &\frac{L_{apo-sym}}{\bar{v}}(P_{apo}(r,t) - P_{sym}(r,t) - RT(C_{apo}(r,t) - C_{sym}(r,t)), \end{aligned} \tag{S39}$$

where $k_{apo}$ and $k_{sym}$ [mol m$^{-1}$ Pa$^{-1}$ s$^{-1}$] are the hydraulic conductivity of the apoplast and symplast, respectively, $L_{apo\text{-}sym}$ [m Pa$^{-1}$ s$^{-1}$] is the permeability of the semi-permeable membrane between the symplast and apoplast that is perfectly impermeable to solutes. Importantly, we note that Eq. S39 accounts for the flux driven by disequilibrium of water potential between the apoplast and symplast and requires tracking of osmotic pressure in both compartments.

Mass conservation requires that changes in the concentration of water molecules, $M_w$ [mol m$^{-3}$] in the apoplast and symplast, are the sum of intra- and inter-compartment fluxes:

$$\frac{\partial M_{w,apo}\left(r,t\right)}{\partial t}=-\frac{1}{r}\frac{\partial\left(rJ_{apo}(r,t)\right)}{\partial r}-\frac{J_{apo-sym}}{l_{apo}},\tag{S40}$$

$$\frac{\partial M_{w,sym}\left(r,t\right)}{\partial t}=-\frac{1}{r}\frac{\partial\left(rJ_{sym}(r,t)\right)}{\partial r}+\frac{J_{apo-sym}}{l_{sym}}\tag{S41}$$

We find the relation between pressure and water content as:

$$\frac{\partial M_{w,apo}}{\partial P_{apo}}=c_{apo},\tag{S42}$$

$$\frac{\partial M_{w,sym}}{\partial P_{sym}}=c_{sym},\tag{S43}$$

where $c_{apo}$ and $c_{sym}$ [mol m$^{-3}$ Pa$^{-1}$] are the hydraulic capacities of the apoplast and symplast. Relative to Eq. S14, here, we require distinct capacitances for each compartment. Finally, the governing equations can be rewritten as:

$$\text{Apoplast:}\frac{\partial P_{apo}\left(r,t\right)}{\partial t}=\frac{k_{apo}}{c_{apo}}\frac{1}{r}\frac{\partial}{\partial r}\left(r\frac{\partial P_{apo}(r,t)}{\partial r}\right)-\frac{L_{apo-sym}}{c_{apo}l_{apo}\bar{v}}\left(\psi_{apo}(r,t)-\psi_{sym}(r,t)\right)\tag{S44}$$

$$\text{Symplast:}\frac{\partial P_{sym}\left(r,t\right)}{\partial t}=\frac{k_{sym}}{c_{sym}}\frac{1}{r}\frac{\partial}{\partial r}\left(r\frac{\partial P_{sym}(r,t)}{\partial r}\right)+\frac{L_{apo-sym}}{c_{sym}l_{sym}\bar{v}}\left(\psi_{apo}(r,t)-\psi_{sym}(r,t)\right).\tag{S45}$$

Prior the wounding event, we assume that the apoplast does not contain any solutes, and that the water potential in the apoplast and symplast is at equilibrium and constant:

$$P_{apo}(r,t<0)=P_{sym}(r,t<0)-RTC_{sym}=P_0.\tag{S46}$$

At the wound site, the boundary condition depends on the solute concentration:

$$P_{sym}(r=l_{sym},t>0)-RTC_{sym}=RTC_{sol}(r=r_{sym},t).\tag{S47}$$

Far from the wound ($r \gg r_{wound}$), we assume no-flux condition:

$$\frac{\partial P_{apo}}{\partial r}\left(r\gg r_{wound},t>0\right)=\frac{\partial P_{sym}}{\partial r}\left(r\gg r_{wound},t>0\right)=0.\tag{S48}$$

We now discuss the effect of solutes related at the wound site on the water transport through the tissue. We assume that that prior to the wounding, the apoplast does not contain any solutes, and that the wounding event results in a release of the symplastic content of wounded cells composed of water and solutes with concentration, $C_{sym}$ into the apoplast such that the solute concentration in the apoplast, $C_{apo}(r,\ t)$ is distributed as follows:

$$C_{apo}(r,t<0)=\begin{cases}C_{sym} & \text{for } 0\le r\le l_{sym}\\ 0 & \text{for } l_{sym}<r\ll r_{wound}\end{cases} \tag{S49}$$

Based on our assessment (assumption 4) that the mass flow in the cell walls that defined the tissue's apoplast is negligible relative to molecular diffusion, we treat the evolution of the solute in the apoplast as being purely diffusive (i.e., no advection). Further assuming no exchange of solutes between the apoplast and the symplast, the governing equation for the diffusion of solutes through the apoplast is:

$$\frac{\partial C_{apo}(r,t)}{\partial t}=D_{sol}\frac{1}{r}\frac{\partial}{\partial r}\left(r\frac{\partial C_{apo}(r,t)}{\partial r}\right), \tag{S50}$$

where $D_{sol}$ [$m^2\ s^{-1}$] is the molecular diffusion coefficient of the solutes. Far from the wound ($r \gg r_{wound}$), we assume no-flux condition:

$$\frac{\partial C_{apo}}{\partial r}\left(r\gg r_{wound},t>0\right)=0. \tag{S51}$$

In the composite model, the plant tissue is considered to have uniform hydraulic, conductivity, $k_t$ and capacity, $c_t$, defined as the area and volume averaged of the individual contributions of the symplast and apoplast:[3]

$$k_t=\frac{l_{sym}}{l_{sym}+l_{apo}}k_{sym}+\frac{l_{apo}}{l_{sym}+l_{apo}}k_{apo}, \tag{S52}$$

$$c_t=\frac{l_{sym}}{l_{sym}+l_{apo}}c_{sym}+\frac{l_{apo}}{l_{sym}+l_{apo}}c_{apo}. \tag{S53}$$

The governing equation for the composite model is:

$$\text{Composite:}\frac{\partial P_t(r,t)}{\partial t}=\frac{k_t}{c_t}\left(\frac{1}{r}\frac{\partial P_t(r,t)}{\partial r}+\frac{\partial^2 P_t(r,t)}{\partial r^2}\right). \tag{S54}$$

We solve numerically the governing equations for the solute in the apoplast (Eq. S50), pressure in the apoplast (S44), symplast (S45) and the tissue of the composite model (S54) using the partial differential equations solver *pdepe* in MATLAB R2022b (Mathworks, Natick, MA, USA). We assume that an appropriate distance far from the wound ($r \gg r_{wound}$) is the vein spacing ($W$), as it denotes the maximal length between the tissues before encountering a vein. We use the following parameters that correspond to a small plant (Table S2): $W = 10^{-3}$ m, $P_0$ = -0.2 MPa, cell osmotic pressure of 0.7 MPa ($C_{sym}$ = 840 mol m$^{-3}$)[16], $D_{sol}$ = $10^{-10}$ m$^2$ s$^{-1}$, $k_{apo}$ = 2.7 ·$10^{-12}$ mol m$^{-1}$ Pa$^{-1}$ s$^{-1}$, $k_{sym}$ = 2.8 ·$10^{-12}$ mol m$^{-1}$ Pa$^{-1}$ s$^{-1}$, $c_{apo}$ = $10^{-3}$ mol m$^{-3}$ Pa$^{-1}$ , $c_{sym}$ = $10^{-2}$ mol m$^{-3}$ Pa$^{-1}$, $L_{apo\text{-}sym}$ =$10^{-13}$ m Pa$^{-1}$ s$^{-1}$, $l_{apo}$ = $10^{-7}$ m, $l_{sym}$ = $10^{-4}$ m.

To compare the predictions from the coupled apoplast-symplast model and the composite model, we compare the total water potential, which keeps track of the equilibrium in the system. Fig. S5a shows a plot of water potential in the apoplast, symplast, and tissue of the composite model 100 seconds after the wounding. Fig. S5b shows the evolution of the error between the apoplast and composite model (black solid), and symplast and composite model (black dashed). Given that the estimated error due to neglecting solute in the apoplast is less than 5% and the experimentally measured properties of the tissue are on the order of $k_t/c_t$ = (1±50%) ×$10^{-10}$ m$^2$s$^{-1}$, the composite model provides an appropriate approximation for the propagation of water potential signals in the tissue.

### B. Effect of solutes on systemic signals

In this section, we discuss the case where the wounding event results in a release of excess of liquid water with a non-negligible concentration of solutes, $C_0$ [mol m$^{-3}$] in the xylem vessels. We lay out the updated governing equations of the framework to account for solutes in the xylem; however, a full exploration of this case is beyond the scope of this study and requires a dedicated study on its own.

Assuming no exchange of solutes between the xylem and the adjacent tissue (assumption 5), and that flow in each xylem vessel follows Poiseuille's law (assumption 3), the transport of solutes in the xylem vessels, $C_{xyl}$ can be described by Taylor-Aris dispersion (See Section S7):

$$\frac{\partial C_{xyl}(x,t)}{\partial t} = D_e \frac{\partial^2 C_{xyl}(x,t)}{\partial x^2} - v_{ves} \frac{\partial C_{xyl}(x,t)}{\partial x}, \quad \text{(S55)}$$

where, $v_{ves}$ [m s$^{-1}$] is the cross-sectional averaged vessel velocity, and $D_e$ [m$^2$ s$^{-1}$] is the effective diffusivity given that depends on the diffusion coefficient of the solutes, $D_{sol}$:

$$D_e = D + \frac{v_{ves}^2 l_{ves}^2}{48D}. \quad \text{(S56)}$$

The vessel velocity is obtained from the xylem pressure as:

$$v_{ves}(x,t) = -k_{ves}\bar{v} \frac{\partial P_{xyl}(x,t)}{\partial x}. \quad \text{(S57)}$$

We assume that prior the wounding, the concentration of solutes in the xylem is negligible:

$$C_{xyl}(x,y,t<0) = 0. \quad \text{(S58)}$$

After the wounding, we assume that the wound site contains excess of liquid water with a non-negligible concentration of solutes, $C_0$:

$$C_{xyl}(x,y,t)\big|_{wound} = C_{0.} \tag{S59}$$

We assume no flux-boundary condition at the xylem-tissue interface:

$$\frac{\partial C_{xyl}}{\partial x}(x,y,t)\bigg|_{xylem-tissue} = \frac{\partial C_{xyl}}{\partial y}(x,y,t)\bigg|_{xylem-tissue} = 0. \tag{S60}$$

Eq. S55 allows us to solve for the solute dynamic in the xylem and obtain the resulting water potential in the xylem as defined in Eq. S1:

$$\psi_{xyl}(x,y,t) = P_{xyl}(x,y,t) - RTC_{xyl}(x,y,t). \tag{S61}$$

To solve for the pressure changes in the xylem and tissue, all the governing equations and boundary conditions remain the same as described in Section S2.B, except for the boundary condition at the wound site for the tissue (Eq. S19). The adjusted boundary condition takes into account that the wound site has non-negligible concentration of solutes:

$$P_t(x,y,t>0)\big|_{wound} - RTC_t = -RTC_{xyl}(x,y,t)\big|_{wound}. \tag{S62}$$

### C. Actively regulated osmotic pressure in tissue

One of the major assumptions in our framework presented in Section S2.A, is that we neglect any actively regulated processes that adjust the solute concentration in the tissue. As a result, we assume that the osmotic pressure in the tissue remains constant in time and space ($\pi_t = RTC_t$ = const). We here lay out the updated governing equations of the framework to account for changes in tissue osmotic potential; however, a full exploration of this case is beyond the scope of this study and requires a dedicated study on its own.

By actively regulated processes that adjust the solute concentration in the tissue, we refer to mechanisms in which the concentration of solutes in individual cells can be expressed as a function, $f$ of various physiological parameters, for example, tissue turgor pressure ($P_t$) and stomatal conductance ($g_s$). Consequently, the solute concentration and, hence, the osmotic pressure in the tissue can be rewritten as:

$$C_t(x,y,z,t) = f(t,P_{t,}g_{s,}...) \rightarrow \pi_t(x,y,z,t) = R\cdot T\cdot f(t,P_{t,}g_{s,}...) \tag{S63}$$

The total tissue water potential then becomes:

$$\psi_t(x,y,t) = P_t(x,y,t) - R\cdot T\cdot f(t,P_t,g_s,...). \tag{S64}$$

Because we no longer can assume that changes in the water potential result exclusively from changes in pressure, we need to rewrite the equation on the water flow in the tissue (Eq. S6). As the water through the symplastic pathways (plasmodesmata-mediated) occurs along gradients of pressure, and the water through the cross-membrane (plasma membrane-mediated, e.g., aquaporin-mediated) occurs through gradient of water potentials due to the presence of semi-permeable membranes, the total water flow in the tissue can be rewritten as:

$$\boldsymbol{J}_t(x,y,t) = -k_{t,PD}\left(\frac{\partial P_t(x,y,t)}{\partial x}\hat{\boldsymbol{x}} + \frac{\partial P_t(x,y,t)}{\partial y}\hat{\boldsymbol{y}}\right) - k_{t,PM}\left(\frac{\partial \psi_t(x,y,t)}{\partial x}\hat{\boldsymbol{x}} + \frac{\partial \psi_t(x,y,t)}{\partial y}\hat{\boldsymbol{y}}\right), \tag{S65}$$

where $k_{t,PD}$ is the hydraulic conductivity of the symplastic pathway, and $k_{t,PM}$ is the hydraulic conductivity cross-membrane pathways.

The second adjustment we need to make is for the case where the semi-permeable membrane between the xylem bundle and the tissue is not perfectly impermeable to solutes, allowing them to move between the xylem and the tissue, and vice versa. For this, we rewrite the transport across the xylem-tissue interface (assumption 5, Eq. S8) as:

$$J_{x-t}(x,t) = \frac{L_{x-t}}{\bar{V}}\left(\left(P_{xyl}(x,y,t) - P_t(x,y,t)\right) - (1-\sigma_s)\left(\pi_{xyl}(x,y,t) - \pi_t(x,y,t)\right)\right)\Big|_{xylem-tissue}, \tag{S66}$$

where $\sigma_s$ [-] is the reflection coefficient of the solute ($0 \leq \sigma_s \leq 1$)[17]. For a semi-permeable membrane that is perfectly impermeable to solutes, we have $\sigma_s = 1$.

With the correction of these two fluxes—water in the tissue (Eq. S65) and water between the xylem and tissue (Eq. 66)—along with the rest of the derivation presented in Section S2.B-C, we can solve for the case where the osmotic pressure is actively regulated. The main challenge in solving this problem lies in identifying and defining the processes that regulate solute adjustment (i.e., defining the function *f* in Eqs. S63 and S64).

### S4. Experimentally measured model inputs

In this section, we provide the experimentally measured physiological values for parameters used in our model and the references from which we accessed them. For the values that were not directly measured in experiments, we summarize their derivation. Table S2 provides a summary of all the physiological values for different plant sizes: small plants (L = $10^{-3}$ – $10^{-2}$ m), e.g., *Arabidopsis thaliana*, medium plants (L = 0.1 – 1 m), e.g., tomato, and large plants (L = 10 – 100 m), e.g., trees.

<u>Xylem bundle diameter and xylem hydraulic conductivity.</u> For a typical radius of xylem vessels ranging from $(5.1 \pm 0.2)\cdot 10^{-6}$ m, (Arabidopsis [18]), $(10\pm 2)\cdot 10^{-6}$ m (tomato [19]), $35 \cdot 10^{-6}$ m (trees [20]), and a xylem

bundle porosity of 80 % , we calculate the xylem hydraulic conductivity (Eq. S5) of the order of $5 \cdot 10^{-5}$, $2 \cdot 10^{-4}$, and $2.5 \cdot 10^{-3}$ mol m$^{-1}$ Pa$^{-1}$ s$^{-1}$ for Arabidopsis, tomato, and trees, respectively.

Xylem hydraulic capacity. Since the xylem vessels are composed of thick lignified cell wall, the value of its hydraulic capacity can be estimated as the inverse of the volumetric modulus of lignified cell wall[3]: $c_{xyl} \sim 1/\left(\bar{v} \cdot \varepsilon_{xyl}\right)$. With the modulus of lignified cell wall of $\varepsilon_{xyl} \sim 1$ GPa,[21] the hydraulic capacity of xylem is the order of $10^{-4}$ mol m$^{-3}$ Pa$^{-1}$.

### S5. Numerical solution of propagation of systematic signals

To obtain the plots shown in Fig. 2, we numerically solve the governing equations for the xylem (Eq. S15) and tissue turgor pressure (Eq. S16) for an arbitrary leaf shape using a finite element analysis software package (COMSOL 6.2, COMSOL Inc., Burlington, MA, USA). We first create a leaf geometry and mesh it using triangular elements as shown in Fig. S1b. Files F1 and F2 provide the geometry and mesh used in this simulation. We then apply initial and boundary conditions (Eqs. S19 - 22) and run a time-dependent simulation. We use the following parameters (Table S2): $L = 10^{-2}$ m, $2W = 0.4 \cdot 10^{-3}$ m, $\kappa_{xyl} = 1$ m$^2$s$^{-1}$, $\kappa_t = 10^{-10}$ m$^2$s$^{-1}$, $P_0 = -0.2$ MPa, $\pi_{xyl} = 0$ MPa, $\pi_t = 0.75$ MPa, $E = 1$ mmol m$^{-2}$ s$^{-1}$, $c_t = 10^{-2}$ mol m$^{-3}$ Pa$^{-1}$, $d_{th} = 200 \cdot 10^{-6}$ m, $h_{stem} = 10^{-8}$ mol m$^{-2}$ Pa$^{-1}$ s$^{-1}$.

Since the xylem time scale is several orders lower than the tissue time scale, $\tau_{\mathrm{xyl}} << \tau_{\mathrm{t}}$ (Eqs. S27 and S28), we first run the simulation for short time (0–100 μs) and fine time steps (0.1 μs) and then perform a second simulation with longer time scales (0–400 s) and coarse time steps (0.5 s). Movie S2 and S3 show time evolution of the xylem and tissue turgor pressure obtained from the simulation. To obtain the plots shown in Fig. 2c and Fig. 2d, we use the simulated 2D solution shown in Movie S2 and S3, respectively, and extract variations along one axis: the x-direction for the xylem bundle and the y-direction for tissue.

### S6. Numerical solutions of leaf swelling

Malone[13] observed that localized scorching of one leaf results in a thickness increase (i.e., leaf swelling) of a neighboring leaf as shown in Fig. 3a. These experiments were performed on a wheat seedling subjected to non-transpiring conditions (high humidity or darkness).

To simulate the predicted increase in leaf thickness with our model and generate the plots in Fig. 3b-c and the simulation shown in Movie S4, we numerically solve the tissue turgor pressure (Eq. S16) across the leaf thickness using *PDEmodel* solver in MATLAB R2022b (Mathworks, Natick, MA, USA). We use the following parameters (Table S2): $\kappa_t = (1 \pm 0.5) \cdot 10^{-10}$ m$^2$s$^{-1}$, $d_{th} = 200 \cdot 10^{-6}$ m, and three value of vein spacing $2W = 0.2 \cdot 10^{-3}$ m, $0.3 \cdot 10^{-3}$ m, $0.4 \cdot 10^{-3}$ m. The MATLAB code is provided in MATLAB Script S1.

**Table S2.** Range of physiological values of model inputs based on experimental measurements.

| Quantity | Symbol | Units | Small plants | Medium plants | Large plants | Ref |
|---|---|---|---|---|---|---|
| Leaf length | $L$ | m | $10^{-3} – 10^{-2}$ | 0.1 – 1 | 10 – 100 | - |
| Vein spacing | $2W$ | m | $(0.4±0.2) \cdot 10^{-3}$ | | | [22] |
| Xylem bundle diameter | $l_{xyl}$ | m | $(1.3±0.1) \cdot 10^{-4}$ | $(2±0.1) \cdot 10^{-4}$ | $2 \cdot 10^{-3}$ | [18–20] |
| Xylem vessel radius | $l_{ves}$ | m | $(5.1±0.2) \cdot 10^{-6}$ | $(10±2) \cdot 10^{-6}$ | $35 \cdot 10^{-6}$ | [18–20] |
| Xylem hydraulic conductivity | $k_{xyl}$ | mol $m^{-1}$ $Pa^{-1}$ $s^{-1}$ | $(5±0.8) \cdot 10^{-5}$ | $(2±1.6) \cdot 10^{-4}$ | $2.5 \cdot 10^{-3}$ | Eq. S5 |
| Xylem vessel conductivity | $k_{ves}$ | mol $m^{-1}$ $Pa^{-1}$ $s^{-1}$ | $(2±0.3) \cdot 10^{-4}$ | $(7.8±0.8) \cdot 10^{-4}$ | $9.6 \cdot 10^{-3}$ | [18–20] |
| Tissue hydraulic conductivity | $k_t$ | mol $m^{-1}$ $Pa^{-1}$ $s^{-1}$ | $10^{-13}$ - $10^{-12}$ | | | $k_{PD}+ k_{PM}$ |
| Tissue apoplast hydraulic conductivity | $k_a$ | mol $m^{-1}$ $Pa^{-1}$ $s^{-1}$ | $2.7 \cdot 10^{-12}$ | | | [23] |
| Tissue symplast hydraulic conductivity | $k_s$ | mol $m^{-1}$ $Pa^{-1}$ $s^{-1}$ | $2.8 \cdot 10^{-12}$ | | | [23] |
| Plasmodesmata hydraulic conductivity | $k_{PD}$ | mol $m^{-1}$ $Pa^{-1}$ $s^{-1}$ | $10^{-13}$ - $10^{-12}$ | | | [24] |
| Plasma membrane hydraulic conductivity | $k_{PM}$ | mol $m^{-1}$ $Pa^{-1}$ $s^{-1}$ | $10^{-14}$ - $10^{-12}$ | | | [25,26] |
| Xylem-tissue permeability | $L_{x-t}$ | m $Pa^{-1}$ $s^{-1}$ | $10^{-12}$ | | | [27,28] |
| Root-stem transport coefficient | $h_{stem}$ | mol $m^{-2}$ $Pa^{-1}$ $s^{-1}$ | $10^{-9}$ - $10^{-8}$ | | | [29] |
| Xylem hydraulic capacity | $c_{xyl}$ | mol $m^{-3}$ $Pa^{-1}$ | $10^{-4}$ | | | [21] |
| Tissue hydraulic capacity | $c_t$ | mol $m^{-3}$ $Pa^{-1}$ | $10^{-2}$ - $10^{-3}$ | | | [27,30] |
| Transpiration flux | $E$ | mol $m^{-2}$ $s^{-1}$ | $(0.1\text{-}10) \cdot 10^{-3}$ | | | [12] |

The geometry considered is depicted in Fig. S6. We assume that the veins (xylem bundles) are arranged periodically with inter-vein spacing of $2W$. Due to this symmetry, we solve only for the periodic domain around one vein. The diameter of the vein is $l_{xyl}$, and the leaf thickness is $d_{th}$. Since the leaf-to-leaf distance in the wheat seedlings used by Malone are in the other of 20 cm, the xylem bundle in that case is expected to approach equilibrium within milliseconds ($\tau_t \cong 40$ ms; Fig. 2e). As a result, we assume that the initial xylem pressure is zero along the entire length of the leaf and solve for the tissue turgor pressure. We assume no-flux conditions on the top and bottom of the leaf and, no-flux boundary condition on the edge of the domain due to symmetry. Movie S4 shows the time evolution simulation of tissue dynamics.

Assuming no exchange of solutes between the xylem and constant osmotic pressure, changes in the water potential are reflected as changes in the turgor pressure in the tissue. We first obtain the leaf volume increase, using the relationship between the leaf bulk elastic modulus, $\varepsilon_t$ [Pa] and the change in turgor pressure as follows:

$$\varepsilon_t = \frac{\Delta P_t(t)}{\frac{\Delta V(t)}{V_0}} \rightarrow \frac{\Delta V(t)}{V_o} = \frac{\Delta P_t(t)}{\varepsilon_t}. \tag{S67}$$

To estimate the leaf bulk elastic modulus, we use the tissue hydraulic capacity:[3]

$$c_t = \frac{1}{\bar{v}(\varepsilon_t + \pi_t)}, \tag{S68}$$

where $\pi_t$ is a typical osmotic pressure of the leaf. Based on our assumption that changes in xylem and tissue volume are negligible in the x-y plane, and experimental observations indicating that the relative inextensibility of the xylem constrains in-plane area changes, any increase in leaf volume during water uptake manifests as an increase in leaf thickness ($\Delta V/V_0 = \Delta d_{th}/d_{th,0}$).[31]

### S7. Numerical solution of propagation of chemical elicitors in xylem vessels

#### A. Methodology

Here, we describe how we obtained the predicted propagation distance of chemical elicitors released at the wound site for comparison with experimental measurements of the propagation of calcium cytosolic signals[32] in a *A. th.* plant (Fig. 4b), and of the propagation of electrical signals[10] in a wheat seedling (Fig. 4c).

The mass flow of water in the xylem vessels from the wound site can carry hypothetical chemical elicitor (i.e., Ricca factor), released at the wound site, into non-wounded leaves by advection-diffusion. The transport of chemical elicitors can be described by Taylor-Aris dispersion as shown in the work of Evans and Morris.[5] For an *A.th.*, the validity of this assumption is supported by the following two conditions: (i) for the expected velocity of a chemical elicitor ($v \cong 500$ μm/s), the diffusion coefficient of the elicitor ($D \cong 100$ μm$^2$s$^{-1}$), and a typical xylem vessel radius ($l_{ves} \cong 5$ μm), the Peclet number, defining the ratio between axial advective and transverse diffusive transport processes ($Pe = v \cdot l_{ves}/D = 25$) is high. (ii) The axial length that an elicitor must travel before it has diffusively sampled all radial locations in the xylem vessel ($L_{TA} = Pe \cdot l_{ves} = 0.125$ mm) is much smaller than the length of the plant ($L \cong 1$ cm). Similarly, for wheat seedling the two conditions are also satisfied: (i) for the expected velocity of a chemical elicitor ($v \sim 0.1$ cm/s), its diffusion coefficient ($D \cong 100$ μm$^2$s$^{-1}$) and a typical xylem vessel radius ($l_{ves} \cong 10$ μm), the Peclet number ($Pe = v \cdot l_{ves}/D = 100$) is high. (ii) The axial length that the elicitors must travel before it has diffusively sampled all radial location of the xylem vessel ($L_{TA} = Pe \cdot l_{ves} = 1$ mm) is much smaller than a typical length of the plant ($L \cong 20$ cm). See this reference[33] for background information on Taylor-Aris dispersion.

The transport of chemical elicitors can be thus described with Taylor-Aris dispersion:

$$\frac{\partial C_{elicitor}(x,t)}{\partial t} = D_e \frac{\partial^2 C_{elicitor}(x,t)}{\partial x^2} - v_{ves} \frac{C_{elicitor}(x,t)}{\partial x} \tag{S69}$$

where $C_{elicitor}$ [mol m$^{-3}$] is the concentration of elicitors, $v_{xyl}$ [m s$^{-1}$] is the cross-sectional average xylem velocity, and $D_e$ is the Taylor-Aris effective diffusivity given by:

$$D_e = D + \frac{v_{ves}^2 l_{ves}^2}{48D} = D(1 + \frac{Pe^2}{48}). \tag{S70}$$

The xylem velocity is obtained from the xylem pressure as:

$$v_{ves}(x,t) = -k_{ves}\overline{v} \frac{\partial P_{xyl}(x,t)}{\partial x}. \tag{S71}$$

The profile of the xylem pressure is obtained from a COMSOL simulation as described below (S6.B-C). For a xylem velocity that varies in time and space, as is the case for a wounding event, Eq. S69 can be solved only numerically. Rather than solving Eq. S69 directly, we calculate the propagation distance of a chemical elicitor by simulating an ensemble 50 trajectories of diffusive tracers in the average flow $v_{xyl}(x, t)$ (i.e., the Lagrangian dynamics of the solutes):

$$x(t) = v_{ves}(x,t) \cdot \Delta t + \Delta x(t), \tag{S72}$$

where $\Delta x(t)$ is a diffusive step implemented during each convective time step with step size chosen as a Gaussian random number with a mean of zero and a standard deviation of the form $\Delta x(t) = \sqrt{2D_e \Delta t}$. We use MATLAB 2022b to run the simulation and choose a time step of $\Delta t = L/(v_{xyl} \cdot 1000)$, which is much shorter than the advective timescale along the xylem vessel. As an initial condition, we impose an instantaneous release of tracers at time $t=0$ at the wound site. To obtain the total displacement, we track the average location of the released tracers as a function of time. It is important to note that axial dispersion $\Delta x_{disp}$ [m] (i.e., dispersive broadening of the concentration of elicitors along the xylem vessel), which in the order of $\Delta x_{disp} \sim \sqrt{D_{eff}\, t}$ , is much smaller relative to the plant length in both small and medium plants, as discussed in the following section S7.B and S7.C.

### B. Parameters for model predictions shown in Fig. 4b

Here, we provide all the parameters used to make model predictions on the propagation of a hypothetical chemical elicitor in *A.th.*, and confront it with experimentally observed calcium signals by Bellandi *et al.*[32] (Fig. 4b). To numerically solve the governing equations for the xylem (Eq. S15) and tissue turgor pressure (Eq. S16), we first use a finite element analysis software package (COMSOL 6.2, COMSOL Inc., Burlington, MA, USA). We create a leaf geometry and mesh it using triangular elements as shown in Fig. S1b. Files F1 and F2 provide the geometry and mesh used in this simulation. We then apply initial and boundary conditions (Eqs. S19 - 22) and run two time-dependent simulation (0–35 s with time step $\Delta t = 0.01$ s): one for a transpiring ($E = 1$ mmol $m^{-2}$ $s^{-1}$), and one for non-transpiring plant ($E = 0$ mmol $m^{-2}$ $s^{-1}$). The rest of the parameters are the same for both simulations (Table S2): $L = 10^{-2}$ m (typical plant length measured by Bellandi *et al.*[32] ), $2W = 0.4\cdot 10^{-3}$ m, $\kappa_{xyl} = 0.5$ $m^2s^{-1}$, $\kappa_t = 10^{-10}$ $m^2s^{-1}$, $P_0 = -0.2$ MPa, $\pi_{xyl} = 0$ MPa, $\pi_t = 0.75$ MPa, $c_t = 10^{-2}$ mol $m^{-3}$ $Pa^{-1}$, $d_{th} = 200 \cdot 10^{-6}$ m, $h_{stem} = 10^{-8}$ mol $m^{-2}$ $Pa^{-1}$ $s^{-1}$. Files F3 and F4 provide the simulated gradient of the xylem pressure in a non-transpiring and a transpiring plant, respectively. Note: in this calculation, we only varied the value of the uniform rate of transpiration, $E$ in order to improve the match between the prediction and experimental measurements. For this adjustment, we overlaid the experimental measurements and predictions and selected the best fit through visual assessment (i.e., by eye).

Using the predicted gradient of the xylem pressure, we then run a Lagrangian dynamics in MATLAB, as described in the previous section S7.A, to simulate the propagation distance of a elicitor with diffusion coefficient $D = 10^{-10}$ $m^2s^{-1}$ in a xylem vessel with radius $l_{ves} = 5 \cdot 10^{-6}$ m , and a range of values of experimentally measured hydraulic conductivities, $k_{ves} = (2\pm0.3) \cdot 10^{-4}$ mol $m^{-1}Pa^{-1}s^{-1}$ (see Table S2). The MATLAB code of the Lagrangian dynamics is provided in Script S2. The resulting values of Taylor-Aris diffusivity (Eq. S70), and axial dispersion, $\Delta x_{disp}$ spanned the following range for transpiring plant: $D_e = 1.3\cdot 10^{-10} - 3.7\cdot 10^{-9}$ $m^2s^{-1}$, $\Delta x_{disp} = 67 - 350$ µm, and for non-transpiring plant: $D_e = 1.1\cdot 10^{-10} - 4.5\cdot 10^{-10}$ $m^2s^{-1}$, $\Delta x_{disp} = 62 - 122$ µm. We note that this axial dispersion remains weak relative to the uncertainty in the propagation distance, as shown in Fig. 4b (blue and grey shaded regions), which can be up to 1000 µm due to variability in the reported available experimental model parameters.

### C. Parameters for model prediction shown in Fig. 4c

Here, we provide all the parameters used to make model predictions on the propagation of a hypothetical chemical elicitor, and confront it with experimentally observed electrical signals by Vodeneev *et al.*[10] (Fig. 4c). We first run a COMSOL simulation to solve for the governing equations for the xylem (Eq. S15) and tissue turgor pressure (Eq. S16) in wheat seedling. Files F5 and F6 provide the geometry and mesh used in this simulation. We then apply initial and boundary conditions (Eqs. S19 - 22) and run two time-dependent simulations (0–150 s with time step $\Delta t = 0.1$ s): one for a transpiring ($E = 1.2$ mmol $m^{-2}$ $s^{-1}$), and one for non-transpiring plant ($E = 0$ mmol $m^{-2}$ $s^{-1}$). The rest of the parameters are the same for both simulations (Table S2): $L = 0.2$ m (typical plant length measured by Vodeneev *et al.*[10]), $2W = 0.4 \cdot 10^{-3}$ m, $\kappa_{xyl} = 1$ $m^2s^{-1}$, $\kappa_t = 10^{-10}$ $m^2s^{-1}$, $P_0 = -0.2$ MPa, $\pi_{xyl} = 0$ MPa, $\pi_t = 0.75$ MPa, $c_t = 10^{-2}$ mol $m^{-3}$ $Pa^{-1}$, $d_{th} = 200 \cdot 10^{-6}$ m, $h_{stem} = 10^{-8}$ mol $m^{-2}$ $Pa^{-1}$ $s^{-1}$. Files F7 and F8 provide the simulated gradient of the xylem pressure in a non-transpiring and a transpiring plant, respectively. Note: similar to the previous case, in this calculation, we only varied the value of the uniform rate of transpiration, $E$ in order to improve the match between the prediction and experimental measurements. For this adjustment, we overlaid the experimental measurements and predictions and selected the best fit through visual assessment (i.e., by eye).

Using the predicted gradient of the xylem pressure, we then run a Lagrangian dynamics in MATLAB to simulate the propagation distance of elicitors with diffusion coefficient $D = 10^{-10}$ $m^2s^{-1}$ in a xylem vessel with radius $l_{ves} = 10 \cdot 10^{-6}$ m , and a range of values of experimentally measured hydraulic conductivities, $k_{ves} = (7.8 \pm 0.8) \cdot 10^{-4}$ mol $m^{-1}Pa^{-1}s^{-1}$ (see Table S2). The MATLAB code of the Lagrangian dynamics is provided in Script S3. The resulting values of Taylor-Aris diffusivity (Eq. S70), and axial dispersion, $\Delta x_{disp}$ spanned the following range for transpiring plant: $D_e = 3.2 \cdot 10^{-9} – 9.9 \cdot 10^{-8}$ $m^2s^{-1}$, $\Delta x_{disp} = 0.69 – 3.9$ mm, and for non-transpiring plant: $D_e = 2.5 \cdot 10^{-9} – 1.6 \cdot 10^{-8}$ $m^2s^{-1}$, $\Delta x_{disp} = 0.67–1.5$ mm. We note again (as in Section S7.C for Fig. 4b) that this axial dispersion remains weak relative to the uncertainty in the propagation distance, as shown in Fig. 4c (blue and grey shaded regions), which can be up to 20 mm due to variability in the reported available experimental model parameters.

### S8. Numerical solution of propagation of local signals

To numerically solve the governing equation for the propagation of the tissue turgor pressure (Eq. S29) in the context of local signaling subjected to the initial (Eq. S30) and boundary conditions (Eqs. S31 and S32), we use the partial differential equations solver *pdepe* in MATLAB R2022b (Mathworks, Natick, MA, USA). Movie S5 shows simulation of the resulting tissue pressure. To obtain the plot in Fig. 5c, we plot the distance at which the tissue turgor pressure is 0.6 MPa. We use the following parameters (Table S2): $\kappa_t = (1 \pm 0.5) \cdot 10^{-10}$ $m^2s^{-1}$, $2W = 10^{-3}$ m, $D = 10^{-10}$ $m^2s^{-1}$, $P_0 = -0.2$ MPa, $\pi_t = 0.75$ MPa, $r_{wound} = 5 \cdot 10^{-6}$ m. The MATLAB code is provided in Script S4.

In the main text (Section: Propagation of local signals and Fig. 5), we explore two hypotheses – molecular diffusion of a chemical elicitor and poroelastic diffusion of a change in turgor pressure – for the propagation of local the local propagation of cytosolic calcium signals reported by Bellandi *et al.*[32] Here, we consider and exclude a third hypothesis: the propagation of elicitors advected with the poroelastic mass flow in the tissue initiated by the wound.

Assuming that there is no solute exchange between cells, and solute distribution in the tissue remains constant (assumption 2), changes in water potential are reflected as changes in tissue pressure. The resulting velocity, $v_t$ [m s$^{-1}$] of the water flow from the wound site to the rest of the tissue as follows:

$$v_t(r,t) = -k_t\bar{v}\frac{\partial P_t(r,t)}{\partial r}. \quad \text{(S73)}$$

Using a similar approach as in Section S7, we employ a numerical simulation to estimate the total displacement a chemical elicitor would travel due to the advective flux resulting from the gradient of the tissue pressure. We choose a time step much smaller than the tissue relaxion scale, $\Delta t = 0.1\text{s} \ll \tau_t \cong 100\text{s}$, and at each time step we displace each of the elicitors a deterministic radial displacement of $v_t \cdot \Delta t$. Fig. S7b shows the predicted displacement from the model. Importantly, we note that the predicted advection with the mass flow in the tissue is ≅ 6 µm, more than one order lower than the experimentally observed displacement ( ≅300 µm; reason for not plotting them on same plot) shown in Fig. 7a, suggesting that advection of an elicitor with the predicted mass flow in the tissue is too weak to contribute significantly to the propagation of cytosolic calcium signals.

While the numerical solution provides a more accurate estimate of the predicted radial propagation of the elicitors, we also use a scaling analysis to obtain a rough estimate of the resulting velocity, $v_t$ as:

$$v_t(r,t) = -k_t\bar{v}\frac{\partial P_t(r,t)}{\partial r} \approx -k_t\bar{v}\frac{0-P_{t,0}}{l_{cell}} = k_t\bar{v}\frac{P_{t,0}}{l_{cell}}, \quad \text{(S74)}$$

where $P_{t,0}$ [Pa] is the initial turgor in the tissue prior the wounding and $l_{cell}$ [m] is a typical length of a cell in the tissue. For a typical cell turgor pressure of $P_{t,0}$ = 1 MPa, cell length of $l_{cell}$ = 10 µm, and the highest measured tissue conductivity of $k_t = 10^{-12}$ mol m$^{-1}$ Pa$^{-1}$ s$^{-1}$ (Table S2), the expected flow velocity is in the order of $v_t$ = 0.18 µm s$^{-1}$. Even if this flow velocity remains constant, the maximal distance that an elicitor would be advected within 200 s, is $d$ = 200s ·0.18 µm s$^{-1}$ = 36 µm, whereas the observed experimentally displacement is in the order of 250 µm (Fig. S7a). Note that the flow velocity is expected to decrease as a function of time due to the tissue relaxation, and therefore, this maximal velocity that we estimated will not be sustained for over 200 s and the expected displacement would be lower than 36 µm as shown in Fig. S7b.

### S9. Propagation of pressure changes as a signaling pathway of systemic signals

Several studies[34–36] have suggested that propagation of pressure changes could serve as an upstream trigger for chemical and electrical systemic signals, with mechanosensitive ion channels as potential candidates for decoding these changes.[37] Although spatiotemporal activation of mechanosensitive channels has not been shown unambiguously to be involved in the propagation of signals, a recent study by Moe-Lange *et al.*[37] has identified a stretch-activated anion channel, MSL10, expressed adjacent to the vasculature, influenced the propagation of wound-induced electrical and calcium signals in *A.th.* plant with typical leaf-to-leaf distance of up to 3 cm: in comparisons of a *msl10* mutant versus wildtype, they observed that the arrival time of an electrical signal in non-wounded leaf was unaffected, but the duration of the signal was shorter in the mutant. Based on this observation, they suggested that systemic signaling requires a combination of chemical ligand perception and activation of mechanosensitive channels.

Based on our predictions of poroelastic propagation in the xylem (Fig. 2c), we hypothesize that pressure changes in the xylem would activate, nearly simultaneously, all mechanosensitive channels along the xylem-tissue interface throughout the entire plant. The timescale for this mode of activation would be defined by the relaxation time in the xylem, $\tau_{xyl} \cong 1$ ms, for a plant with total leaf-to-leaf distance, $L \cong$ 3 cm. Therefore, we suggest that pressure changes alone cannot trigger the chemical and electrical systemic signals that have been observed to propagate progressively over many seconds.[32,37,38] Instead, pressure changes could act as primers for tissue to propagate signals with slower dynamics, potentially driven by mass flow of chemical elicitors as discussed in Section *Driving mechanisms for mass flow of chemical elicitors in xylem* (main text) and Section S7. Additionally, we note that the rapid arrival of this pressure signal may directly trigger responses that have not been observable with the reporters used to date or due to insufficient temporal resolution of the measurement techniques employed.

### S10. Potential experiments to dissect signaling pathways in local signals

In the main text (Section *Propagation of local signals,* Fig. 5), we explore two scenarios in which local signals are triggered by: (i) molecular diffusion of an elicitor released at the wound site that can bind to ligand-gated channels (Fig. 5a(i)), and (ii) propagation of pressure change that could induce mechanotransduction of cellular responses (Fig. 5a(ii)). As noted in the main text, molecular diffusion (Eq. 3 in Fig. 5a, scenario (i)), and poroelastic diffusion of pressure (Eq. 4 in Fig. 5a, scenario (ii)) are described by the same form of diffusion equation with similar diffusivities ($D \cong \kappa_t \cong 10^{-10}$ m$^2$ s$^{-1}$ ) such that the rate at which pressure changes propagate is expected to be similar to that of diffusion of small molecules.[39] Consequently, the readout of calcium signals could be explained by either the diffusive spread of chemical elicitors or the propagation of pressure changes as shown in Fig. 5c, or a combination of both processes.

Bellandi *et al.*[32] showed that the dynamics of local calcium signals remain unaffected in plants with plasmodesmal blockage achieved by callose deposition, suggesting that calcium signals are most likely triggered by an apoplastic diffusion of a chemical elicitor (scenario (i)). In order to determine whether the closure of plasmodesmata could affect the propagation of pressure changes, we need to estimate how much the symplastic pathways contribute to the overall flow of water in the tissue. While hydraulic conductivity of plasma membrane-mediated pathways can be experimentally measured in protoplasts[25], the conductivity of plasmodesmata-mediated pathways is typically estimated theoretically, through calculations[40]. The predicted conductivity of plasmodesmata depends on their aperture,[41] thus reducing this aperture by blocking the plasmodesmata should lead to a decrease of the hydraulic conductivity of plasmodesmata-mediated pathways. However, given the uncertainty regarding which pathway that dominates water transport in the overall tissue—plasma membrane-mediated or plasmodesmata-mediated— due to the lack of experimental measurements, it remains an open question as to how the propagation of pressure changes are affected by plasmodesmal blockage. Potential experiments that could tease apart whether local calcium signals are triggered by pressure changes, or diffusion of a chemical elicitor include: (i) simultaneous measurement of calcium signals and tissue pressure, which is possible using a recently developed reporter of apoplastic water potential in the mesophyll, AquaDust;[42] and (ii) measurement of calcium dynamics in mutants that lack target mechanical transducers such as MSL channels, which would be defective in perceiving changes in pressure.

### S11. Reversal of the direction of transpiration-driven flow in wounded leaves

In a non-wounded transpiring plant, the soil serves as the only source of water for the transpiration-driven flow through the xylem that moves from the stem to the petiole and on to the tips of the leaves, as shown in Fig. S8a. Following the wounding of a leaf tip, the release of water from ruptured cells into the open ends of disrupted xylem vessels allows the wound site to serve as a second source of water for mass flows driven by both poroelastic relaxations and continued transpiration (Fig. S8b). To understand the effect of the new source created by the wound, consider the simplified representations of the hydraulic architecture in Fig. S8. In the unwounded plant (Fig. S8a), well-watered soil ($\psi_{soil}$ = 0 MPa) passes through a resistance representing the root and stem ($R_{\mathrm{stem}}$) to feed transpiration in the leaves. Upon wounding (Fig. S8b), the wound site becomes a second source ($\psi_{wound}$ = 0 MPa) that is connected to the stem through a hydraulic resistance, $R_{\mathrm{leaf}}$. Considering only the component of xylem flow driven by transpiration in the unwounded leaf (Fig. S8b), we see that it is pulled through an effective resistance defined by $R_{\mathrm{stem}}$ and $R_{\mathrm{leaf}}$ in parallel. For paths in parallel, the larger flow rate passes through the branch with the lower resistance. For the two paths of interest, we estimate (Table S2):

$$\frac{R_{leaf}}{R_{stem}} = \frac{L/2 \cdot k_{xyl}}{1/h_{stem}} = \frac{L \cdot h_{stem}}{2 \cdot k_{xyl}} \sim 10^{-6}. \qquad \text{(S75)}$$

Therefore, we expect the wet wound site becomes the overwhelmingly dominant source of water after wounding, with flow from the wound toward the stem in the xylem of the wounded leaf (Fig. S8b), opposite of the direction in which it flows in the unwounded case (Fig. S8a).

The reversal of the transpiration stream in the wounded leaf continues until all the water released at the wound site is exhausted, at which point the tension in the xylem is re-established, and the transpiration stream resumes its normal direction (from stem to leaf tip), as discussed in Section S12. During this reversal, both water influx at the wound site and transpiration contribute to the total mass flow in transpiring plants, while in non-transpiring plants, mass flow relies solely on water influx at the wound site. Consequently, the magnitude of mass flow in transpiring plants is higher than that in non-transpiring plants, as shown in Fig. S8b.

### S12. Difference between wet and dry wounds

We define a wet wound in which the wounded leaf is immersed in water, and the water at the wound site is not depleted during the signal propagation. In contrast, in a dry wound the wounded leaf is exposed to air, and the only water released at the wound site comes from the symplastic content of wounded cells. Experimental observations have shown that in wet wounds, the observed signal dynamics (e.g., calcium and electrical signals) propagate into distal leaves, whereas in dry wounds, the signals are only propagated within the wounded leaf (i.e., it did not reach the stem or other leaves).[32,43] Here, we use our framework to model these two type of wounds, and make predictions of the resulting propagation distances of chemical elicitors released at the wound site (Fig. S9).

In a wet wound, we assume that the wound site has excess of available liquid water as described in Eq. S9:

$$P_{xyl}(x,y,t>0)\big|_{wound} = P_t(x,y,t>0)\big|_{wound} - \pi_t = 0. \qquad \text{(S76)}$$

In a dry wound, the wound site initially has available excess liquid water for a certain time, $\tau_{wet}$ [s], until the water is depleted:

$$P_{xyl}(x,y,\tau_{wet}>t>0)\big|_{wound} = P_t(x,y,\tau_{wet}>t>0)\big|_{wound} - \pi_t = 0. \qquad \text{(S77)}$$

After this period, the water in the xylem pins to the border pit membrane as shown in Fig. S9a, and the boundary condition becomes a non-flux:

$$\nabla P_{xyl}(x,y,t>\tau_{wet})\cdot\hat{\boldsymbol{n}}_{wound} = \nabla P_t(x,y,,t>\tau_{wet})\cdot\hat{\boldsymbol{n}}_{wound} = 0. \qquad \text{(S78)}$$

where $\hat{\boldsymbol{n}}_{wound}$ is the normal vector at the wound.

Following the same procedure and model parameters described in Section S7.A-B for a small herbaceous plant, we obtain the maximal predicted propagation distance of chemical elicitors released at the wound site. The differences between the dry and wet wounds lies in their boundary conditions are: for the wet wound, we apply Eq. S76 continuously, while for the dry wound, we apply Eq. 77 for an initial period, $t$ = 0-5 s to simulate the period required to evacuate a small amount of water released by wounding and then transition to applying Eq. S78 for $t$ > 5 s. For this representation of the dry wound, the 5 seconds of flow corresponds to a total volume of 12 nL. Since typical epidermal cells have a volume of 100 to 500 pL,[44,45] this volume corresponds to wounding of approximately 20 to 100 cells. As shown in Fig. S9b, for a dry wound, the maximal distance chemical elicitors propagate is approximately 0.3 cm, remaining within the wounded leaf (which is 0.4 cm long). In contrast, a wet wound that has more water available at the wound site, sustains the mass flow for a longer duration, thus enabling the elicitors to propagate further, reaching non-wounded leaves.

### S13. Movie captions

**Movie S1. Simulation of the initial water potential.** Numerical prediction of a typical steady-state of xylem and tissue turgor pressure prior to the wounding. The simulation was obtained using COMSOL with the following parameters that correspond to a small herbaceous plant (e.g., *Arabidopsis thaliana*) (Table S2): $L = 10^{-2}$ m, $2W$=$0.4\cdot10^{-3}$ m, $\kappa_{xyl}$ =1 $m^2s^{-1}$, $\kappa_t$ =$10^{-10}$ $m^2s^{-1}$, $P_0$ = -0.2 MPa, $\pi_{xyl}$ = 0 MPa, $\pi_t$ = 0.75 MPa, E = 1 mmol $m^{-2}$ $s^{-1}$, $c_t = 10^{-2}$ mol $m^{-3}$ $Pa^{-1}$, $d_{th} = 200 \cdot 10^{-6}$ m, $h_{stem} = 10^{-8}$ mol $m^{-2}$ $Pa^{-1}$ $s^{-1}$.

**Movie S2. Propagation of xylem pressure.** Numerical prediction of the evolution of xylem pressure with the poroelastic model (colorbar; blue – higher pressure, red – lower pressure). Upon wounding, the released water and tension propagates rapidly through the xylem such that the pressure along this path approaches equilibrium (dark blue) within a relaxation timescale, that is in the order of hundreds of microseconds for a small plant. The simulation was obtained using COMSOL with the following parameters that correspond to a small herbaceous plant (e.g., *Arabidopsis thaliana*) (Table S2): $L$ = $10^{-2}$ m, $2W$=$0.4\cdot10^{-3}$ m, $\kappa_{xyl}$ =1 $m^2s^{-1}$, $\kappa_t$ =$10^{-10}$ $m^2s^{-1}$, $P_0$ = -0.2 MPa, $\pi_{xyl}$ = 0 MPa, $\pi_t$ = 0.75 MPa, E = 1 mmol $m^{-2}$ $s^{-1}$, $c_t = 10^{-2}$ mol $m^{-3}$ $Pa^{-1}$, $d_{th} = 200 \cdot 10^{-6}$ m, $h_{stem} = 10^{-8}$ mol $m^{-2}$ $Pa^{-1}$ $s^{-1}$.

**Movie S3. Propagation of tissue pressure.** Numerical prediction of the evolution of tissue pressure with the poroelastic model. Assuming that the osmotic pressure in the tissue remains constant ($\pi_t$=const), the water efflux from the xylem along its entire length and into the tissue induces an increase of tissue pressure over time as indicated by the colorbar (blue – higher pressure, red – lower pressure). The tissue pressure approaches equilibrium within its relaxation timescale that is in the order of hundreds of seconds. As in Movie S3, the simulation was obtained using COMSOL with the following parameters that correspond to a small herbaceous plant (e.g., *Arabidopsis thaliana*) (Table S2): $L = 10^{-2}$ m, $2W$=$0.4\cdot10^{-3}$ m, $\kappa_{xyl}$ =1 $m^2s^{-1}$, $\kappa_t$ =$10^{-10}$ $m^2s^{-1}$, $P_0$ = -0.2 MPa, $\pi_{xyl}$ = 0 MPa, $\pi_t$ = 0.75 MPa, E = 1 mmol $m^{-2}$ $s^{-1}$, $c_t = 10^{-2}$ mol $m^{-3}$ $Pa^{-1}$, $d_{th} = 200 \cdot 10^{-6}$ m, $h_{stem} = 10^{-8}$ mol $m^{-2}$ $Pa^{-1}$ $s^{-1}$.

**Movie S4. Wound-induced leaf swelling.** Numerical prediction of the normalized leaf thickness for three different vein spacings following a wounding event. The simulation was obtained using MATLAB with the following parameters that correspond to a medium plant (e.g., wheat seedling) (Table S2): $\kappa_t = (1\pm0.5) \cdot 10^{-10}\ m^2s^{-1}$, $d_{th} = 200\cdot10^{-6}$ m, and three value of vein spacing $2W = 0.2\cdot10^{-3}$ m, $0.3\cdot10^{-3}$ m, $0.4\cdot10^{-3}$ m. The MATLAB code used to obtain this simulation is provided in MATLAB Script S1.

**Movie S5. Propagation of local tissue pressure.** Numerical prediction of the evolution of the local tissue pressure upon wounding a cell in the middle of the domain. The simulation was obtained using the partial differential equations solver *pdepe* in MATLAB with the following parameters that correspond to a non-transpiring small herbaceous plant (e.g., *Arabidopsis thaliana*) (Table S2): $\kappa_t = (1\pm0.5) \times 10^{-10}\ m^2s^{-1}$, $2W = 10^{-3}$ m, $D = 10^{-10}\ m^2s^{-1}$, $P_0 = -0.2$ MPa, $\pi_t = 0.75$ MPa. The MATLAB code used to obtain this simulation is provided in MATLAB Script S4.

### S14. MATLAB scripts and COMSOL files

**MATLAB Script S1.** This script provides the code used to simulate the predicted increase in leaf thickness (Section S6) and generate the plots in Fig. 3b-c and Movie S4.

**MATLAB Script S2.** This script provides the code used to obtain model predictions on the propagation of a hypothetical chemical elicitor in *A.th* (Section S7.B) and generate the plots in Fig. 4b.

**MATLAB Script S3.** This script provides the code used to obtain model predictions on the propagation of a hypothetical chemical elicitor in wheat seedling (Section S7.C) and generate the plots in Fig. 4c.

**MATLAB Script S4.** This script provides the code used to solve the propagation of local signals (Section S8) and generate Movie S5.

**COMSOL Files F1.** This file contains the geometry used in the COMSOL simulation for solving the dynamic of xylem and tissue pressure in *A.th* (Section S7.B).

**COMSOL Files F2.** This file contains the mesh used in the COMSOL simulation for solving the dynamic of xylem and tissue pressure in *A.th* (Section S7.B).

**COMSOL Files F3.** This file contains the simulated gradient of the xylem pressure in a non-transpiring *A.th* (Section S7.B).

**COMSOL Files F4.** This file contains the simulated gradient of the xylem pressure in transpiring *A.th* (Section S7.B).

**COMSOL Files F5.** This file contains the geometry used in the COMSOL simulation for solving the dynamic of xylem and tissue pressure in wheat seedling (Section S7.C).

**COMSOL Files F6.** This file contains the mesh used in the COMSOL simulation for solving the dynamic of xylem and tissue pressure in wheat seedling (Section S7.C).

**COMSOL Files F7.** This file contains the simulated gradient of the xylem pressure in a non-transpiring wheat seedling (Section S7.C).

**COMSOL Files F8.** This file contains the simulated gradient of the xylem pressure in transpiring wheat seedling (Section S7.C).

### S15. Supplementary figures

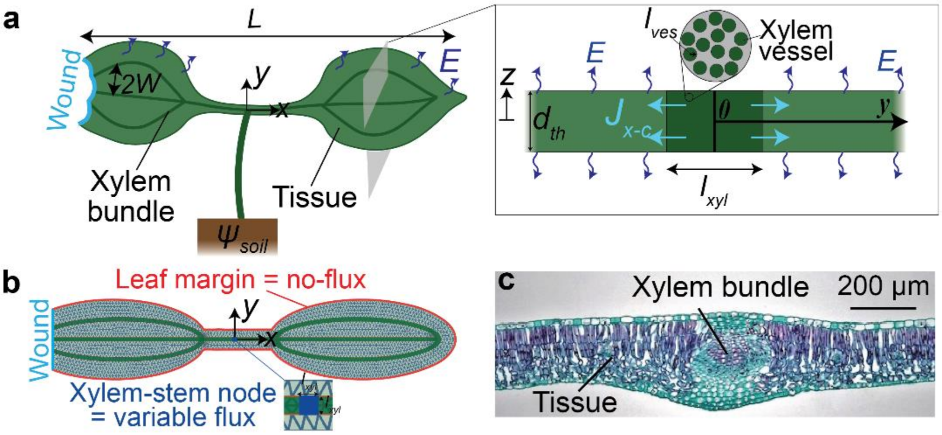

**Fig. S1.** Model geometry and parameters. **a,** Schematic diagram showing the general structure of a thin leaf composed of xylem bundle (dark green) and tissue (light green). As illustrated in the expanded view (right), the xylem bundle and tissue architectures are treated as single fluid-carrying plates with uniform properties through their thickness, $d_{th}$. The tip of one of the leaves is wounded by cutting, burning, or squeezing the vasculature. **b**, Meshed representation of the wounded plant used to numerically solve the resulting gradients of the water potential using COMSOL (See Section S5 for details on the simulation). **c,** Microscope image of a cross-section of a leaf showing a thin plate anatomy composed of tissue and a xylem vascular bundle (adapted from[46]).

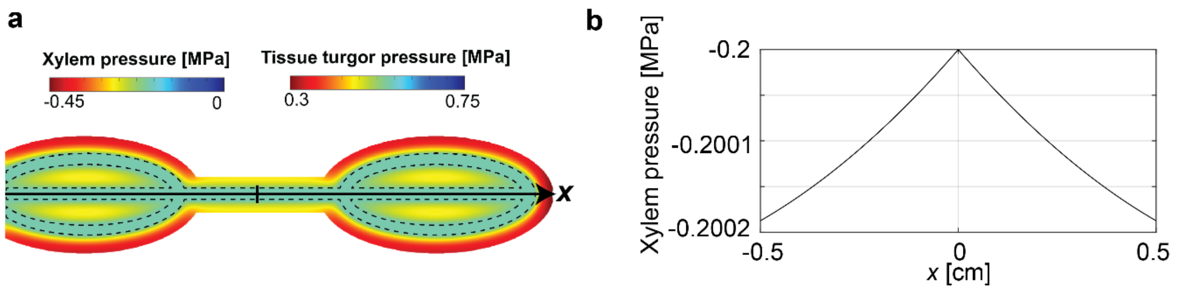

**Fig. S2.** Calculation of initial distribution of xylem and tissue turgor pressure with uniform transpiration. **a**, Numerical prediction of a typical steady-state pressure in the xylem (Eq. S15) and tissue (Eq. S16) prior to the wounding. The region of the xylem bundles is marked with dashed lines. The constant transpiration creates a water potential gradient, driving water from the xylem to the surrounding tissue. Movie S1 shows the time evolution of the pressure dynamics. **b**, Plot of the axial (along black solid line) variation of the xylem pressure in the midvein showing that the xylem pressure drops from the stem toward the tip of both leaves. To obtain these predictions, we ran a COMSOL simulation for 4000 s with the following parameters that correspond to a small plant (Table S2): $L = 10^{-2}$ m, $2W$=0.4 $\cdot 10^{-3}$ m, $\kappa_{xyl}$=1 $m^2s^{-1}$, $\kappa_t = 10^{-10}$ $m^2s^{-1}$, $P_0$ = -0.2 MPa, $\pi_t$ = 0.75 MPa, $\pi_{xyl}$ = 0 MPa, $E$ = 1 mmol $m^{-2}$ $s^{-1}$, $c_t = 10^{-2}$ mol $m^{-3}$ $Pa^{-1}$, $d_{th}$ = 200 · $10^{-6}$, $h_{stem} = 10^{-8}$ mol $m^{-2}$ $Pa^{-1}$ $s^{-1}$.

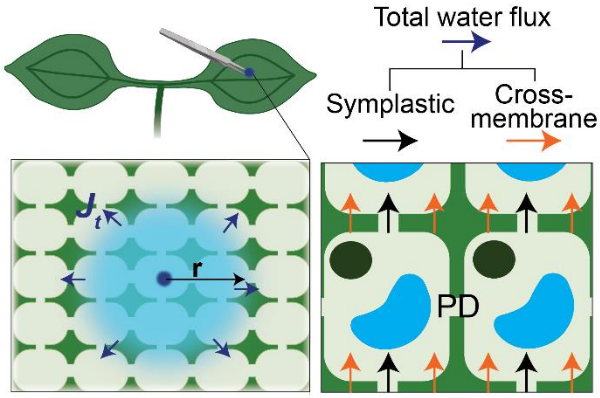

**Fig. S3.** Model of local signals. A schematic diagram showing a local wounding of cells far from the vasculature resulting in propagation of a water potential signal that spreads radially from the wound site. The cells are assumed to be single fluid-carrying regions with uniform tissue properties. The water moves across the tissue predominantly via two pathways: cross-membrane (i.e., plasma membrane-mediated) and symplastic pathways.

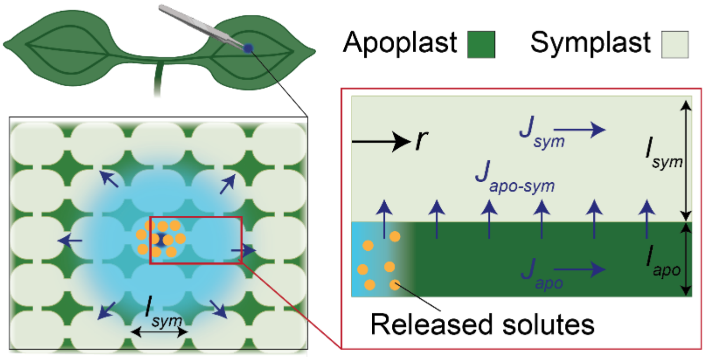

**Fig. S4.** Schematic diagram illustrating the structure of tissue composed of symplast (light green) and adjacent apoplasts (dark green). After a wounding event, the symplastic water containing solutes (orange circles) from wounded cells, is released into the apoplast. From there, water can propagate through the apoplast, move from the apoplast to the symplast, and travel within the symplast. However, solutes can only move through the apoplast because the apoplast-symplast interface is impermeable to solutes.

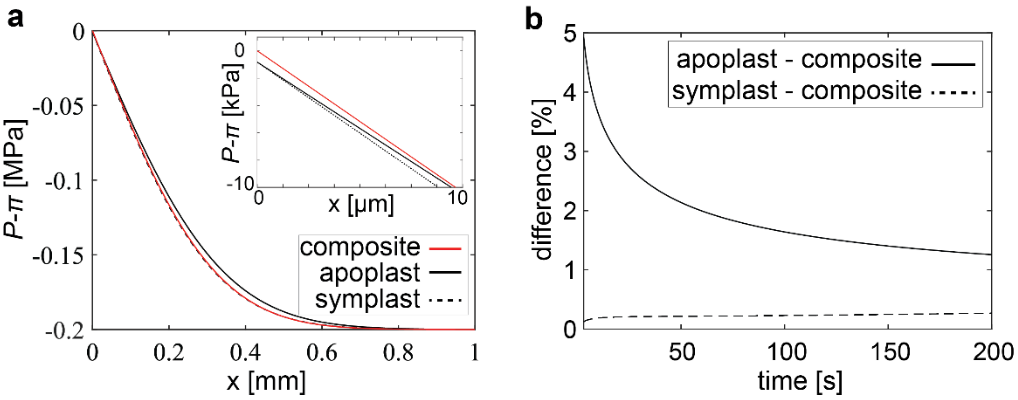

**Fig. S5.** Comparison between composite model and a coupled apoplast-symplast. **a**, Plot of the water potential in the apoplast (black solid line, Eq. S44), symplast (black dashed line, Eq. S45) and composite model (red solid line, Eq. S54) at t = 100 s after the wounding event. We here plot the water potential ($\psi = P - \pi$), which is a function of the turgor ($P$) and osmotic ($\pi$) pressure, as it keeps track of the equilibrium in the system. The inset shows the water potential near the wound site. The water potential in the apoplast and symplast is lower due to the presence of solutes released at the wound site. **b,** Difference between the predicted values of the water potential in the apoplast and composite model (solid line), and symplast and composite model (dashed line) as function of time. The difference between the apoplast and composite is calculated as $diff = \text{abs}(\psi_{apo} - \psi_t)/\psi_t$, and the error between the symplast and composite is calculated as $diff = \text{abs}(\psi_{sym} - \psi_t)/\psi_t$.

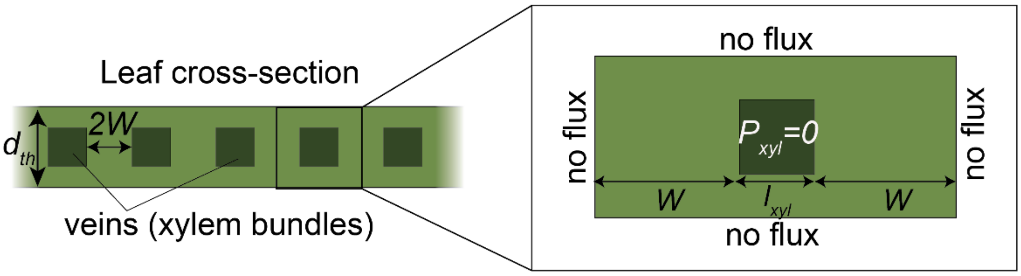

**Fig. S6.** Geometry used for numerically solving the dynamics of turgor pressure (Eq. S16) in the tissue (light green) assuming that the pressure in the xylem (dark green) is zero. We use MATLAB to solve for one domain around a single vein (See Section S6). Movie S4 shows the resulting time evolution simulation of the tissue pressure dynamics for different vein spacing.

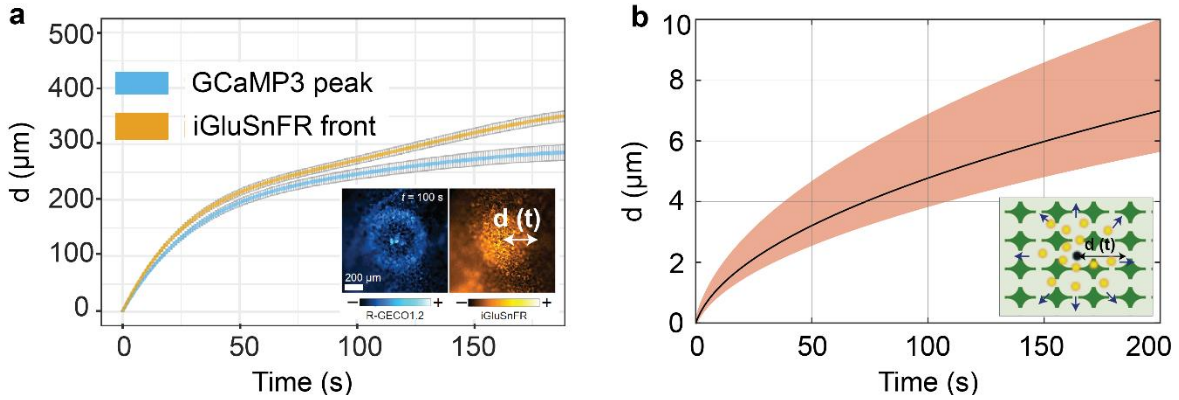

**Fig. S7. a,** Experimentally observed cytosolic calcium signal and apoplastic glutamate (i.e., potential elicitor) signals observed in independent reporter lines: GCaMP3 (reports cytosolic calcium) and iGluSnFR (reports apoplastic glutamate), where *d* is the radial distance of propagation of the signals (adapted from Bellandi *et al.*[32]). The inset shows fluorescence image for calcium and glutamate response in R-GECO1.2 (reports cytosolic calcium) and iGluSnFR (reports apoplastic glutamate) dual reporter. **b**, Time evolution of the radial propagation of an elicitor advected with the flux resulting from the gradient of the tissue pressure. The predicted propagation distance is one order of magnitude lower than the experimentally observed one, suggesting that the observed calcium signals are most likely not triggered by the advection of an elicitor. The model considers a range of values of the tissue poroelastic diffusivity based on experimental measurements ($\kappa_t$ = (1±0.5) $\cdot 10^{-10}$ $m^2s^{-1}$), where the black line corresponds to a value of $\kappa_t = 10^{-10}$ $m^2s^{-1}$, initial turgor pressure of tissue of $P_{t,0}$ = 1 MPa, and a highest measured tissue conductivity of $k_t = 10^{-12}$ mol $m^{-1}$ $Pa^{-1}$ $s^{-1}$ (Table S2).

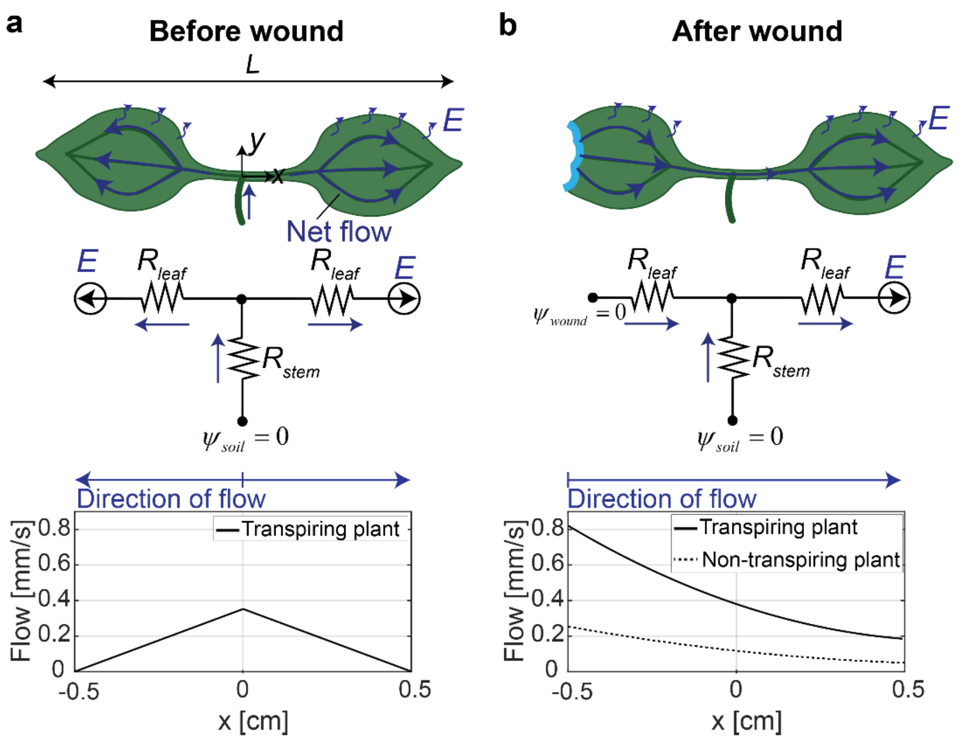

**Fig. S8.** Mass flow of water in xylem in a small herbaceous plant with $L$ = 1 cm (e.g., *A.th.*). **a,** Steady-state transpiration-driven flow prior to wounding in a plant with transpiration flux $E$ = 1 mmol m$^{-2}$ s$^{-1}$. The transpiration-driven flow through the xylem moves from the stem to the leaf tips, with flow magnitude decreasing along the way due to constant transpiration flux, $E$. **b**, Mass flow in wounded plants 1 s after wounding in a transpiring (solid line, $E$ = 1 mmol m$^{-2}$ s$^{-1}$), and non-transpiring plant (dashed line). After wounding, due to the released water at the wound site, the mass flow in the wounded leaf is reversed: moving from the wound toward the stem. In non-transpiring plants (dashed line), the flow is driven solely by poroelastic relaxation of the tissue. In transpiring plants (solid line), this flow is reinforced by transpiration, which reverses its direction in the wounded leaf. In the unwounded plant (a), well-watered soil ($\psi_{soil}$ = 0 MPa) feeds transpiration through root and stem resistance ($R_{stem}$). After wounding (b), the wound site ($\psi_{wound}$ = 0 MPa) adds a second source, connected to the stem via leaf resistance ($R_{leaf}$). To obtain the mass flow predictions, we used COMSOL to simulate the gradient in xylem pressure and the resulting mass flows, as described by Eq. S71 (see Section S7.B for model parameters).

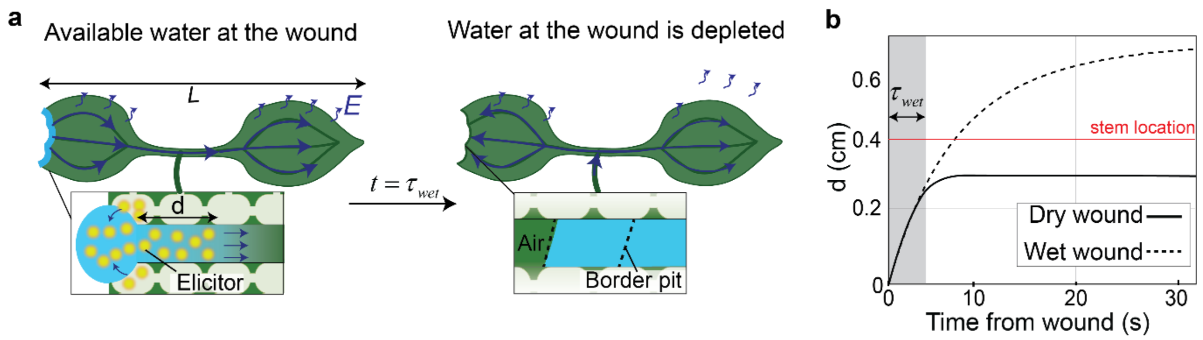

**Fig. S9.** Difference in the signal propagation between a wet wound (i.e., immersing the wounded leaf in water) and dry wound (i.e., exposing the wounded leaf to air). **a,** In a dry wound, the water released at the wound site from the ruptured cells is drawn into the xylem bundles and fed to the tissue, initiating a mass flow towards non-wounded leaves carrying chemical elicitors released at the wound site. Upon water depletion at the wound site (after $t = \tau_{wet}$), the water in the xylem is pined at the border pith membrane, the xylem tension is re-established and the net flow follows the transpiration stream, moving from the stem-to-petiole-to-leaf tip. **b**, Model prediction of the propagation distance, $d$ [m] of a chemical elicitor in a small herbaceous plant with $L = 1$ cm (e.g., *A.th.*) and transpiration flux $E = 1$ mmol m$^{-2}$ s$^{-1}$, for a dry (solid lines) and wet (dashed line) wound. The red horizontal line indicates the location of the stem (i.e., for d < 0.4 cm, the elicitor remains in the wounded leaf, and for d ≥ 0.4 cm, the elicitor reaches the non-wounded leaf). We assume that the water at the wound site is depleted after $\tau_{wet} = 5$s. To obtain these predictions, we followed the same procedure and model parameters to obtain the plot in Fig. 4b (See Section S7.B for model parameters). For the wet wound, we applied the boundary condition in Eq. S76, while for the dry wound, we applied the boundary conditions in Eqs. S77 and S78.
